# T cell self-reactivity during thymic development dictates the timing of positive selection

**DOI:** 10.1101/2021.01.18.427079

**Authors:** Lydia K. Lutes, Zoë Steier, Laura L. McIntyre, Shraddha Pandey, James Kaminski, Ashley R. Hoover, Silvia Ariotti, Aaron Streets, Nir Yosef, Ellen A. Robey

## Abstract

Functional tuning of T cells based on their degree of self-reactivity is established during positive selection in the thymus, although how positive selection differs for thymocytes with relatively low versus high self-reactivity is unclear. In addition, preselection thymocytes are highly sensitive to low-affinity ligands, but the mechanism underlying their enhanced TCR sensitivity is not fully understood. Here we show that murine thymocytes with low self-reactivity experience briefer TCR signals and complete positive selection more slowly than those with high self-reactivity. Additionally, we provide evidence that cells with low self-reactivity retain a preselection gene expression signature as they mature, including genes previously implicated in modulating TCR sensitivity and a novel group of ion channel genes. Our results imply that thymocytes with low self-reactivity down-regulate TCR sensitivity more slowly during positive selection, and suggest that modulation of membrane ion channel function may play a role in regulating TCR tuning throughout development.

**Impact Statement:** Developing T cells whose TCRs have relatively low reactivity experience very brief TCR signaling events, delayed positive selection, and do not fully down regulate their TCR sensitivity as they mature.

## Introduction

T cell fate is dictated by the strength of the interaction between the T cell receptor (TCR) and self-peptide major histocompatibility complexes (self-pMHCs) presented on thymic-resident antigen presenting cells (APCs). While thymocytes whose TCRs interact too weakly or too strongly with self-pMHC undergo death by neglect or negative selection respectively, thymocytes whose TCRs react moderately with self-pMHC can survive and give rise to mature CD4+ or CD8+ single positive (SP) T cells. Naïve CD4 and CD8 T cells that emerge from the thymus were initially thought to be relatively homogeneous, with differences emerging only after antigenic stimulation. However, recent studies have shown that positively selected T cells span a range of self-reactivities, and that positive selection results in functional tuning of T cells based on their degree of self-reactivity(Azzam et al. 1998; Fulton et al. 2015; Persaud et al. 2014; Weber et al. 2012; Mandl et al. 2013). On Naïve T cells, surface expression levels of the glycoprotein CD5 serve as a reliable marker for self-reactivity(Azzam et al. 2001; Azzam et al. 1998; Hogquist and Jameson 2014). CD5 also negatively regulates TCR signals, thereby serving as part of the tuning apparatus(Azzam et al. 2001; Azzam et al. 1998; Hogquist and Jameson 2014; Persaud et al. 2014; Tarakhovsky et al. 1995). T cells with relatively high self-reactivity (CD5^high^) expand more rapidly upon priming and exhibit greater sensitivity to cytokines(Fulton et al. 2015; Persaud et al. 2014; Weber et al. 2012; Gascoigne and Palmer 2011), whereas T cells with relatively low self-reactivity (CD5^low^) survive better in the absence of cytokines and exhibit stronger proximal TCR signals upon stimulation(Persaud et al. 2014; Palmer et al. 2011; K. Smith et al. 2001; Cho et al. 2016). How these distinct functional properties are imprinted during positive selection in the thymus remains unknown.

Positive selection is a multi-step process in which CD4+CD8+ double positive (DP) thymocytes experience TCR signals, migrate from the thymic cortex to the medulla, and eventually downregulate either CD4 or CD8 co-receptor. During this process, thymocytes remain motile and experience serial, transient TCR signaling events, to eventually reach a threshold of signaling in order to complete positive selection(Au-Yeung et al. 2014; Ross et al. 2014; Melichar et al. 2013). Thymocytes bearing MHC-II specific TCRs take 1-2 days to complete positive selection and become mature CD4 SP thymocytes, whereas thymocytes bearing MHC-I specific TCRs give rise to CD8 SP thymocytes about 2-4 days after the initiation of positive selection(Saini et al. 2010; Lucas, Vasseur, and Penit 1993; Kurd and Robey 2016). Interestingly, some studies report that CD8 T cells continue to emerge more than one week after the beginning of positive selection(Saini et al. 2010; Lucas, Vasseur, and Penit 1993). It remains unknown why some thymocytes move expeditiously through positive selection, while others require more time to mature.

TCR signaling leads to a rise in cytosolic calcium concentration, and calcium flux during positive selection induces a migratory pause that may prolong the interaction between thymocyte and APC, thus promoting positive selection(Bhakta and Lewis 2005; Bhakta, Oh, and Lewis 2005; Melichar et al. 2013). Calcium flux in thymocytes is also important for activation of the calcium-dependent phosphatase calcineurin and the downstream transcription factor NFAT, both of which are required for normal positive selection(Gallo et al. 2007; Canté-Barrett, Winslow, and Crabtree 2007; Macian 2005; Neilson et al. 2004; Hernandez, Newton, and Walsh 2010; Oh-hora and Rao 2008). Although the initial calcium flux following TCR stimulation comes from the release of calcium from the endoplasmic reticulum (ER) stores, calcium influx from the extracellular space is needed for prolonged calcium signals(Oh-hora and Rao 2008). During mature T cell activation, depletion of ER calcium stores triggers the influx of calcium via calcium release-activated calcium (CRAC) channels. However, components of the CRAC channels are not required for positive selection(Oh-Hora 2009; Gwack et al. 2008; Vig and Kinet 2009), and the ion channels required for calcium influx during positive selection remain unknown.

Just after completing TCRαβ gene rearrangements and prior to positive selection, DP thymocytes (termed preselection DP) are highly sensitive to low affinity ligands(Davey et al. 1998; Lucas et al. 1999; Gaud, Lesourne, and Love 2018). This property is thought to enhance the early phases of positive selection, but must be down-regulated to prevent mature T cells from responding inappropriately to self(Hogquist and Jameson 2014). A number of molecules have been identified which are involved in modulating TCR signaling in DP thymocytes, and which are downregulated during positive selection. These include the microRNA miR-181a: which enhances TCR signaling by downregulating a set of protein phosphatases(Li et al. 2007), Tespa1: a protein that enhances calcium release from the ER(Lyu et al. 2019; Liang et al. 2017), components of a voltage gated sodium channel (VGSC): that prolongs calcium flux via an unknown mechanism(Lo, Donermeyer, and Allen 2012), and Themis: a TCR associated protein with controversial function(Kakugawa et al. 2009; Patrick et al. 2009; Lesourne et al. 2009; Johnson et al. 2009; Fu et al. 2009; Choi, Cornall, et al. 2017; Fu et al. 2013; Choi, Warzecha, et al. 2017; Mehta et al. 2018). While a diverse set of potential players have been identified, a clear understanding of the mechanisms that render preselection DP thymocytes sensitive to low affinity self-ligands remains elusive.

To understand how self-reactivity shapes TCR tuning during thymic development, we compared the positive selection of thymocytes with MHC-I-restricted TCRs of low, intermediate, or high self-reactivity. We show that positive selecting signals in thymocytes with low self-reactivity occur with a more transient calcium flux, and without a pronounced migratory pause. In addition, thymocytes with low self-reactivity proceed through positive selection more slowly, with a substantial fraction taking over a week to complete positive selection. Furthermore, we show that cells with lower self-reactivity retain higher expression of a “preselection” gene expression program as they mature. This gene set including genes known to modulate TCR signals and a novel set of ion channels genes. These results indicate that T cell tuning during thymic development occurs via developmentally layered sets of T cell tuning genes, maximizing the TCR signaling potential of thymocytes with low self-reactivity throughout their development.

## Results

### Altered TCR signaling kinetics during positive selection of thymocytes with low self-reactivity

To investigate how self-reactivity impacts CD8 SP development, we used TCR transgenic (TCRtg) mice expressing MHC-I-restricted TCRs of low, intermediate, or high self-reactivity. Mice expressing rearranged TCR α and β transgenes from a T cell clone (TG6) with low self-reactivity, exhibit a strong skewing to the CD8 lineage even in a *Rag2* sufficient background (**Fig. S1**). This is indicative of robust positive selection, and is in contrast to the HY TCR transgenic model where low self-reactivity is accompanied by extensive endogenous receptor rearrangement and weak skewing to the CD8 lineage(Azzam et al. 2001; Kisielow et al. 1988). Compared to CD8SP from wild type mice, CD8SP thymocytes from TG6 mice express low levels of CD5, a reliable surface marker of T cell self-reactivity (**Fig. 1a, b**)(Azzam et al. 1998; Mandl et al. 2013; Fulton et al. 2015; Persaud et al. 2014; Azzam et al. 2001). CD8SP thymocytes from TG6 mice also have relatively low expression of Nur77, a marker of recent TCR signaling that also correlates with self-reactivity (**Fig. 1c**)(Fulton et al. 2015). In contrast, CD8SP from two well-characterized TCRtg models, OT-1 and F5, have high and intermediate levels of these markers respectively (**Fig. 1a-c**)(Fulton et al. 2015; Cho et al. 2016; Dong et al. 2017; Kieper, Burghardt, and Surh 2004; Palmer et al. 2011; Ge et al. 2004). The relative levels of CD5 and Nur77 in lymph node T cells of TCRtg mice follows a similar pattern, except that expression of Nur77 in F5 thymocytes decreases somewhat in T cells relative to thymocytes **(Fig. 1d)**. This may reflect weaker recognition of the peripheral versus thymic self-peptides by the F5 TCR. Thus, these 3 TCRtg models span the range of self-reactivity for CD8 T cell positive selection, with TG6 providing an example of low self-reactivity coupled with relatively efficient positive selection.

**Figure 1:**
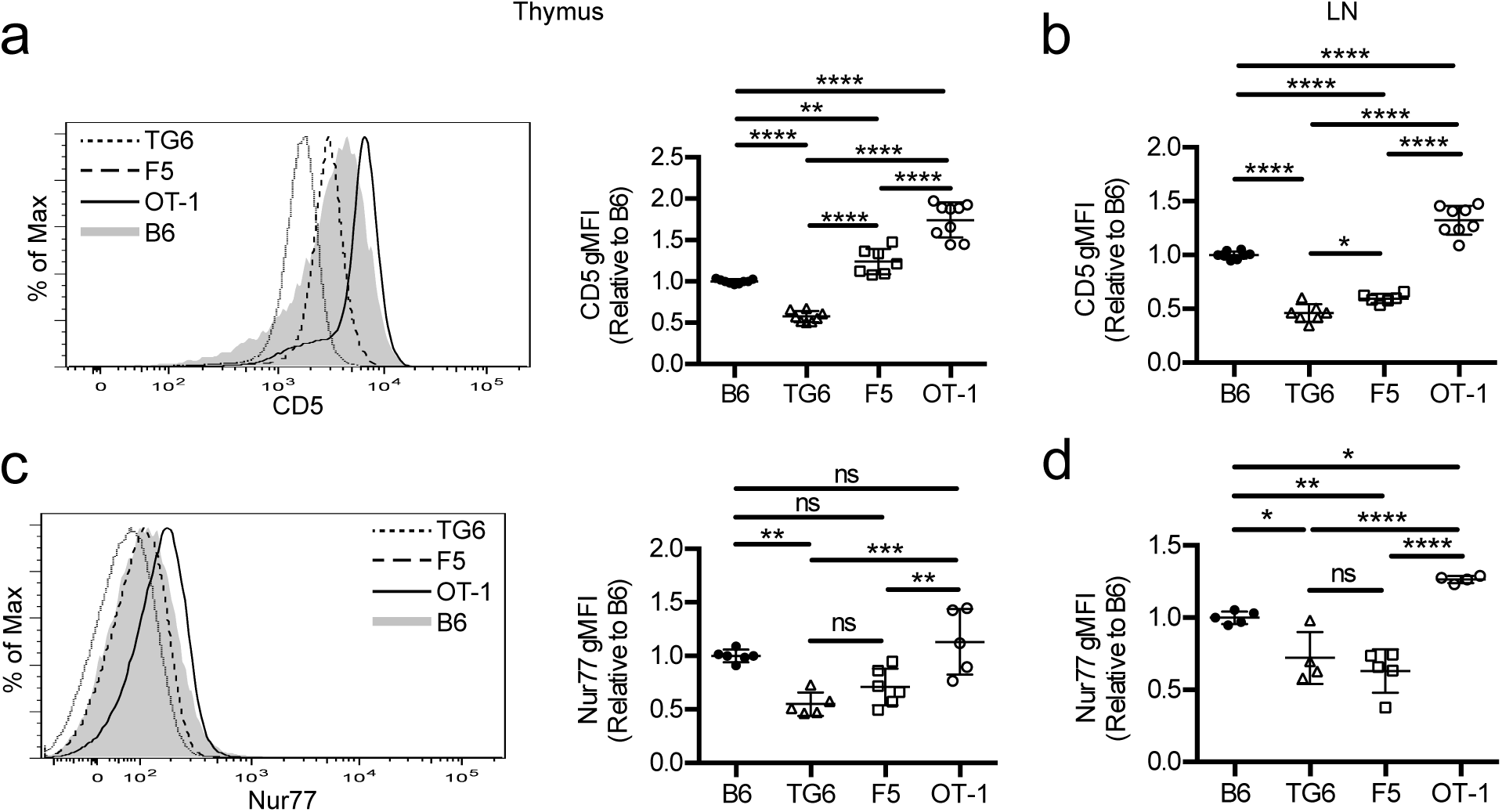
TG6, F5, and OT-1 CD8SP cells express CD5 and Nur77 across the polyclonal spectrum. CD8SP cells were harvested from the thymus and lymph nodes of wild-type (B6), TG6, F5, and OT-1 TCRtg mice and analyzed by flow cytometry. (a and b) Representative (left) and quantified by geometric mean fluorescence intensity (right) CD5 surface expression on (a) CD8SP thymocytes and (b) CD8SP lymph node cells. (c and d) Representative (left) and quantified (right) intracellular Nurr77 expression gated on (c) CD8SP thymocytes and (d) CD8SP lymph node cells. Data are presented as average ± SD and analyzed using an ordinary one-way ANOVA followed by a Tukey’s multiple comparisons (*P <0.05, **P <0.01, ***P <0.001, ****P <0.0001) All data are compiled from 3 or more experiments.

Previous time-lapse studies of OT-1 thymocytes undergoing positive selection *in situ* revealed serial TCR signals lasting about 4 minutes and associated with a migratory pause(Ross et al. 2014; Melichar et al. 2013). To investigate how thymocyte self-reactivity impacts TCR signaling kinetics during positive selection, we compared the pattern of calcium changes and speed for TG6 or OT-1 thymocytes at 3 and 6 hours after the initiation of positive selection. We isolated thymocytes from TCRtg mice on a non-selecting background (herein referred to as preselection thymocytes), labeled the cells with the ratiometric calcium indicator dye Indo-1LR, overlaid the labeled cells onto selecting or non-selecting thymic slices, and imaged using two-photon, time-lapse microscopy as previously described (**Fig. 2a**)(Bhakta, Oh, and Lewis 2005; Ross et al. 2014; Melichar et al. 2013; Dzhagalov et al. 2012; Ross et al. 2015). For both TG6 and OT-1, the frequency of time points with elevated calcium was greater on selecting, compared to nonselecting thymic slices, a difference that was statistically significant at the 6 hour timepoint (**Fig. 2b** and **c**). This confirmed that calcium flux under these conditions is dependent on the presence of positive selecting ligands.

**Figure 2:**
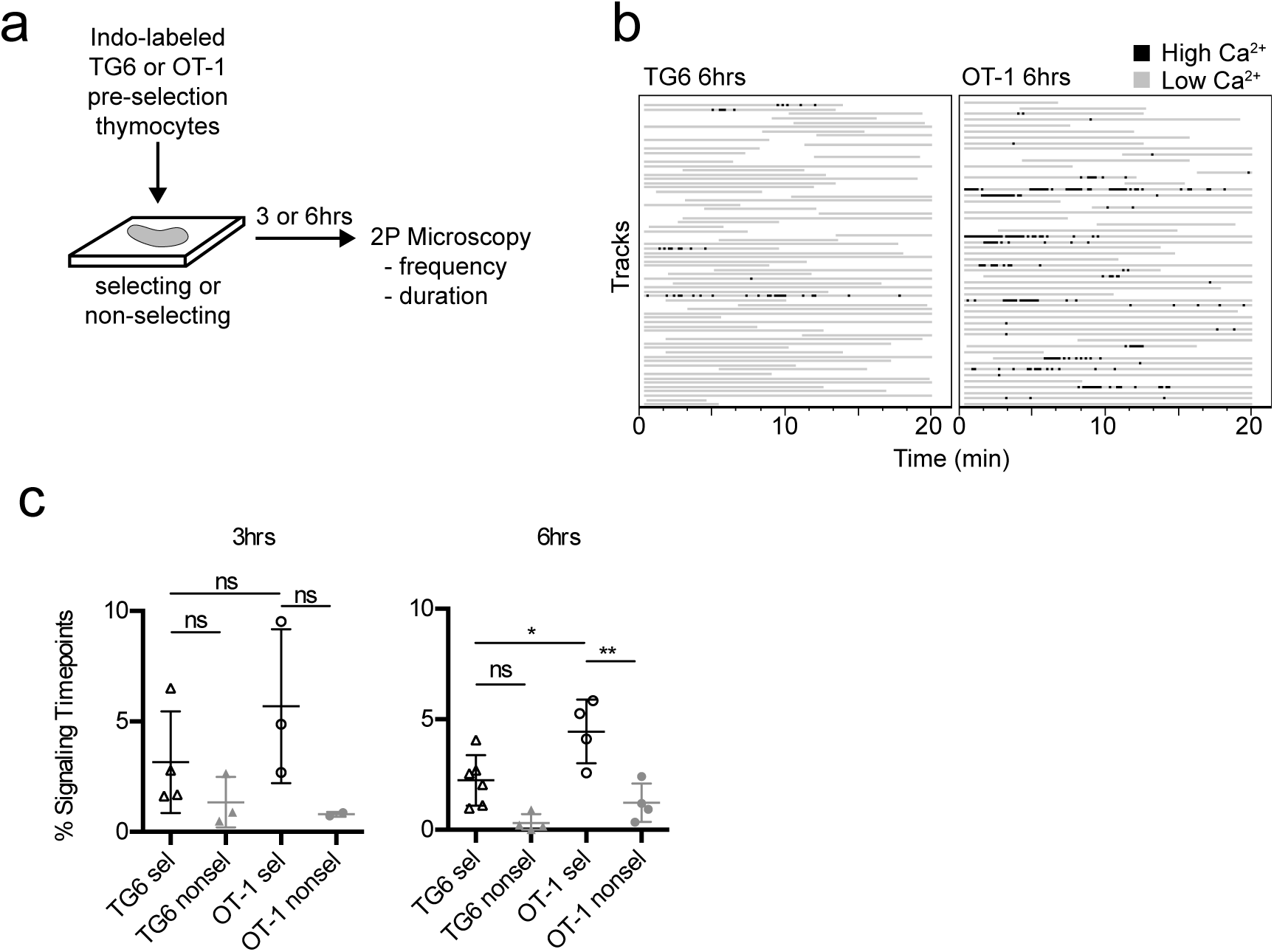
TG6 thymocytes experience less frequent TCR signaling than OT-1 during positive selection. (a) Experimental schematic. (b) Individual tracks (1 track = 1 cell; each track is a horizontal line) over time from TG6 (left) and OT-1 (right) representative runs (movies) 6 hours post addition to the slice. Grey indicates low Ca^2+^ timepoints; black indicates high Ca^2+^ timepoints. (c) Percentage of signaling timepoints (high Ca^2+^ ratio) out of total timepoints for each run. Each dot represents a single run. All data are complied from 2 or more experiments, except OT-1 3 hours data, which is from one experiment. Preselection thymocyte populations were obtained from TG6 Rag2KO H2b mice and irradiated B2MKO mice reconstituted with OT-1 Rag2KO bone marrow. Data are presented as average ± SD and analyzed using an ordinary one-way ANOVA followed by a Tukey’s multiple comparisons (*P<0.05, **P<0.01).

In line with their lower self-reactivity, TG6 cells in positively selecting slices spent significantly less time signaling than did OT-1 cells (**Fig. 2b**, **2c**, **S2** and **Table S1**). To further explore this difference, we identified individual signaling events from multiple imaging runs, aligned the events based on the first timepoint with elevated calcium, and averaged the calcium and speed over time as previously described(Melichar et al. 2013) (see Materials and Methods section). At both 3 and 6 hours after addition to the slice, the average signaling event for TG6 thymocytes is about one minute shorter than OT-1 (**Fig. 3a** and **b**; **Video S1** and **S2**). Moreover, OT-1 thymocytes pause during signaling events, as reflected in a significant decrease in speed at signaling compared to non-signaling timepoints (**Fig. 3c** and **S3**). In contrast, TG6 thymocytes have a much less pronounced pause during signaling events, particularly at the 6-hour time point (**Fig. 3c** and **S3).** Although reduced signal duration contributes to the reduced time TG6 cells spend signaling, this difference could also be due to differences in TCR signaling frequency. Indeed, the frequency of signaling events for TG6 thymocytes was approximately half that for OT-1 thymocytes (1 signaling event every 2 hours for TG6 thymocytes, versus 1 signaling event every 75 minutes for OT-1: **Table S2**). Despite differences in signaling frequency, the percentage of signaling tracks per run was comparable between the two transgenics (**Fig. S3**). Thus, low self-reactivity is associated with less pronounced TCR-induced pausing and a reduction in both the duration and frequency of TCR signals during positive selection.

**Figure 3:**
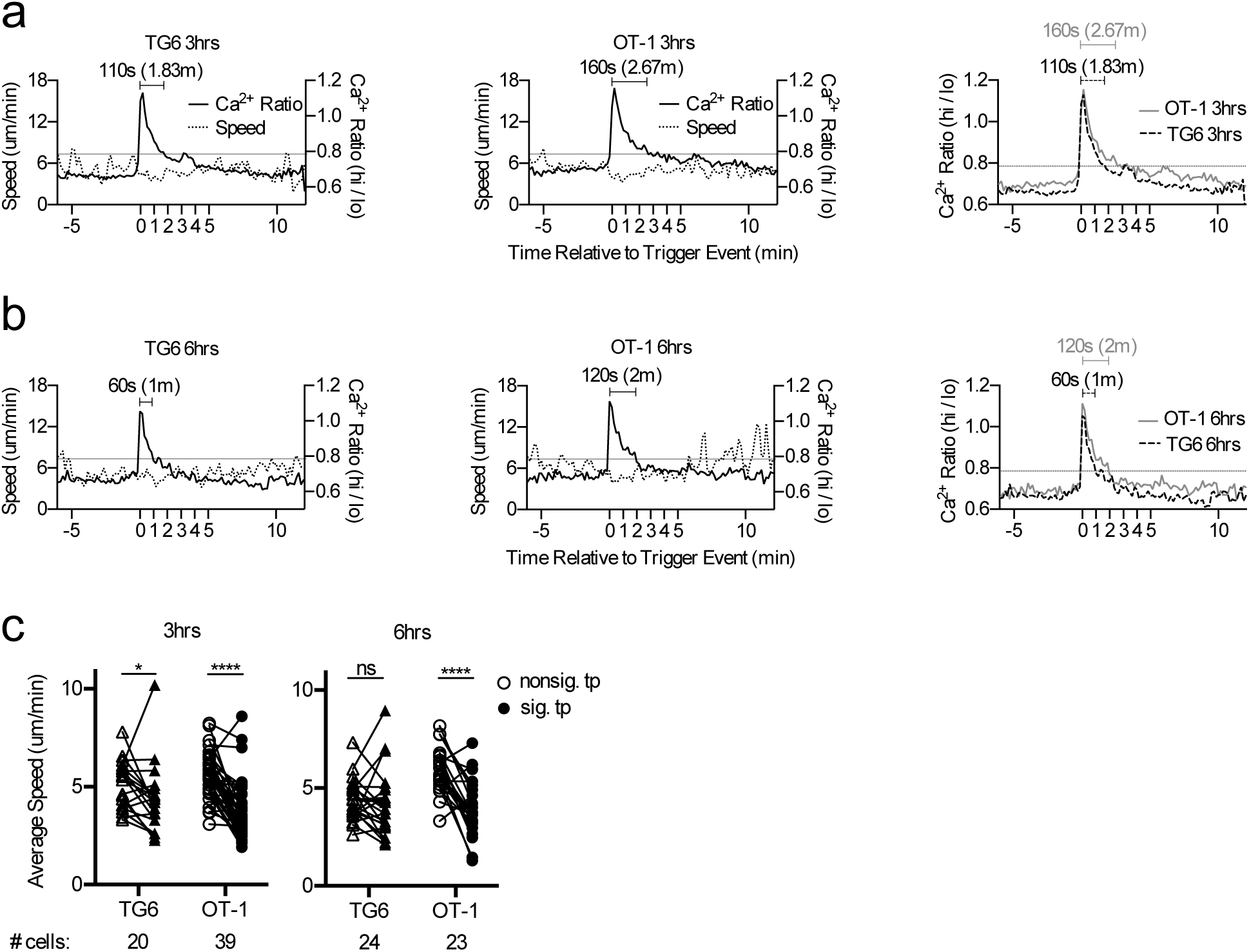
TG6 thymocytes experience TCR signals of shorter duration and a less pronounced signal-associated migratory pause compared to OT-1 during positive selection. Average calcium (Ca^2+^) ratio and speed of signaling tracks (cells) for the indicated conditions, aligned by signaling trigger event (time=0), over time, displayed separately for each TCRtg (left and middle) and overlaid (right). (a) Average aligned Ca^2+^ ratio and speed 3 hours post addition to the slice (TG6 n=27 cells, OT-1 n=37 cells). (b) Average aligned Ca^2+^ ratio and speed 6 hours post addition to the slice (TG6 n=25 cells, OT-1 n=23 cells). Straight horizontal line at Ca^2+^ ratio=0.785 indicates the Ca^2+^ ratio of the end of the event following the trigger event. (c) Average speed of signaling (sig) and nonsignaling (nonsig) timepoints for tracks with at least one signaling event. In each column, a dot represents a single track. Between columns, lines connect data from the same track. All data are compiled from two or more experiments, except OT-1 3hrs data, which is from one experiment. Preselection thymocyte populations were obtained from TG6 Rag2KO H2b mice and irradiated B2MKO mice reconstituted with OT-1 Rag2KO bone marrow. Data are presented as average ± SD and analyzed using a two-way ANOVA with Sidak’s multiple comparisons (*P<0.05, ****P<0.0001).

### Self-reactivity correlates with time to complete positive selection

Reduced TCR signals observed in TG6 thymocytes could mean that thymocytes with low self-reactivity need more time to accumulate sufficient signals to complete positive selection(Au-Yeung et al. 2014). To address this possibility, we examined the timing of different stages of positive selection using thymic tissue slice cultures(Ross et al. 2015). We previously showed that MHC-I specific preselection DP thymocytes undergo a synchronous wave of maturation after introduction into positive selecting thymic tissue slices. This includes transient upregulation of the TCR activation marker CD69, followed by a switch in chemokine receptor expression from cortical (CXCR4) to medullary (CCR7) in DP thymocytes, and eventually CD4 down-regulation(Ross et al. 2014; Melichar et al. 2013). To examine the impact of self-reactivity on the timing of these landmarks in positive selection, we overlaid preselection TG6, F5, or OT-1 thymocytes onto selecting and non-selecting thymic slices, and analyzed progress through positive selection by flow cytometry after 2, 24, 48, 72 and 96 hours of thymic slice culture. Slice-developed TG6, F5, or OT-1 CD8SPs exhibit the same relative CD5 levels as their *in vivo* counterparts (**Fig. 4a**), indicating that thymic slices culture provides a good model for examining the impact of self-reactivity on positive selection. In all three models, thymocytes exhibited a transient increase in CD69 expression, however cells with lower self-reactivity reached peak CD69 expression later than cells with higher self-reactivity (**Fig. 4b**). In addition, the switch in cortical to medullary chemokine receptor expression (CXCR4-CCR7+) and the appearance of CD8SP were delayed in thymocytes with lower self-reactivity (**Fig. 4c, d**, **S4**, and **S5**). Thus, the timing of all three major landmarks during positive selection is inversely correlated with self reactivity, implying that cells with lower self-reactivity progress more slowly through positive selection.

**Figure 4:**
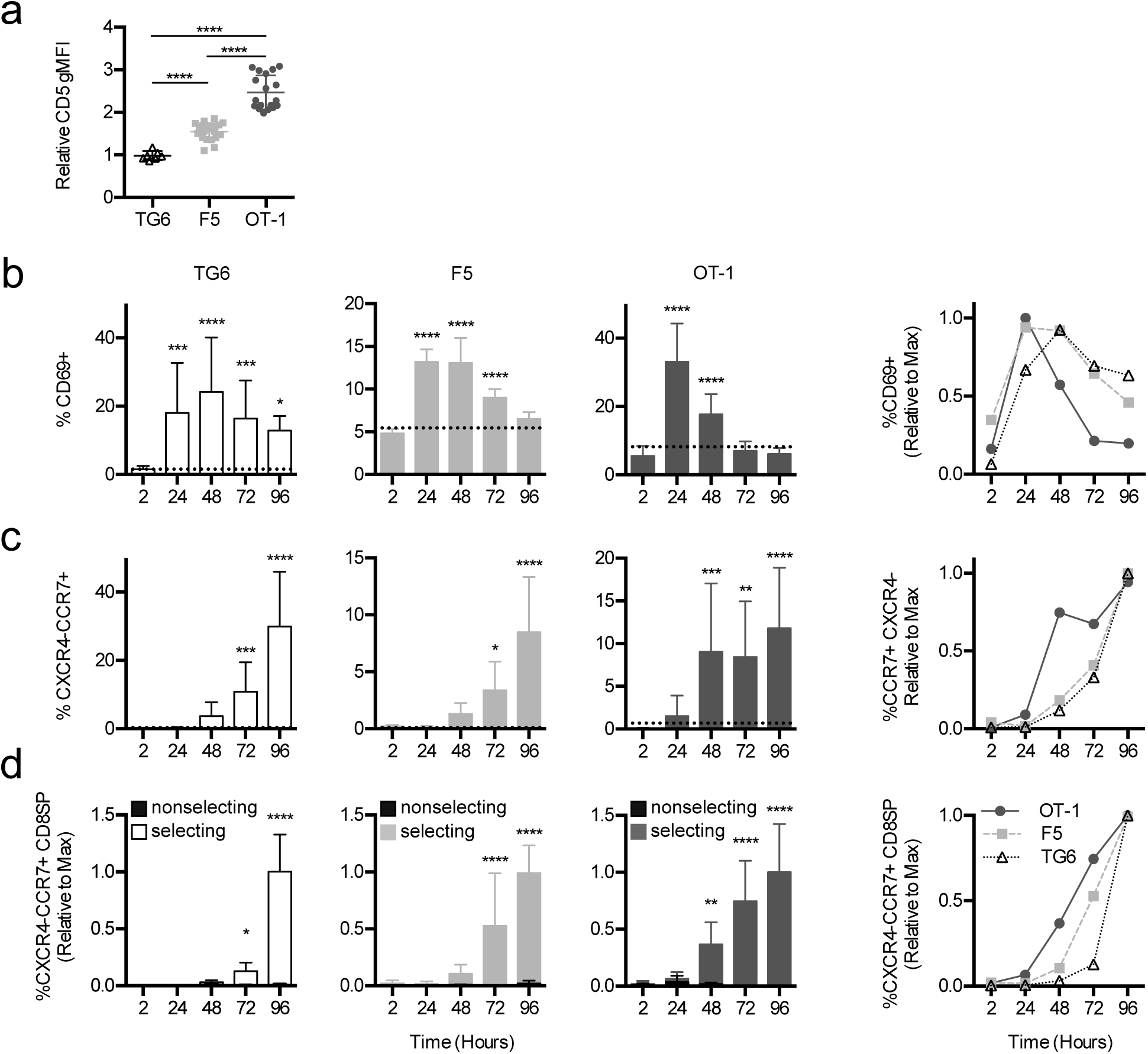
Thymocytes with lower self-reactivity exhibit delayed in situ positive selection kinetics relative to those with high self-reactivity. (a) Expression of CD5 on CXCR4-CCR7+ CD4-CD8+ TG6 (96h), F5 (72h and 96h), and OT-1 (72h and 96h) donor cells developed in thymic slices. Values for each experiment are normalized to the average CD5 expression of the CXCR4-CCR7+ CD4-CD8+ polyclonal slice resident (SR) population at the same timepoints. (b-d) Preselection TG6 (white bars, triangle), F5 (grey bars, square), or OT-1 (dark grey bars, circle) thymocytes were overlaid onto selecting or non-selecting thymic slices, harvested at 2, 24, 48, 72 or 96 hours post thymocyte overlay, and analyzed using flow cytometry. Left three graphs show individual values for each transgenic. Horizontal lines in a and b indicate the average value for non-selecting slices. Right line graph shows the average for all three transgenics overlaid. (b) Percent of CD69+ cells within the CD4+CD8+ and CD4-CD8+ populations. (c) Percentage of CXCR4-CCR7+ cells within CD4+CD8+ and CD4-CD8+ populations. (d) Percentage of CXCR4-CCR7+ CD4-CD8+ cells out of the donor population. Values for each experiment are normalized to the average at the timepoint with the maximum CD8SP development (72h or 96h). For (a) data are presented as average ± SD and analyzed using an ordinary one-way ANOVA with Tukey’s multiple comparisons (****P <0.000). For (b-d) data are presented as average ± SD and analyzed using an ordinary one-way ANOVA with Dunnett’s multiple comparisons (**P <0.01, ***P <0.001, and ****P <0.0001) comparing to 2 hour timepoint. All data are compiled from 3 or more experiments. Preselection thymocyte populations were obtained from TG6tg Rag2KO H2b mice, irradiated B2MKO mice reconstituted with F5tg Rag1KO bone marrow, and OT-1tg Rag2KO B2MKO mice. See also Figure S4 and S5.

Mature CD8SP thymocytes appear around the end of gestation in mice and gradually accumulate after birth. If thymocytes with low self-reactivity proceed more slowly through positive selection, they may also appear more slowly during the neonatal period. To test this prediction, we performed flow cytometry on the thymus and spleen from TG6, F5, and OT-1 mice at 7, 14, and 21 days after birth. Consistent with thymic slice data, CD8SP thymocytes accumulate slowly with age in TG6 mice, with the percent of CD8SP remaining significantly below maximal adult levels at 21 days after birth (**Fig. 5a** and **b**). In contrast, in OT-1 mice, CD8SP thymocytes reach their maximal levels at 7 days after birth, and F5 mice show an intermediate rate of accumulation (**Fig. 5a** and **b**). In addition, TG6 mice were slowest to accumulate CXCR4-CCR7+ CD4+CD8+DP and CD8SP thymocytes (**Fig. 5c**), and exhibited delayed appearance of CD8SP cells in the spleen (**Fig. S6**). These results are consistent with results from thymic slice cultures, and provide *in vivo* confirmation that the time to complete positive selection correlates inversely with thymocyte self-reactivity.

**Figure 5:**
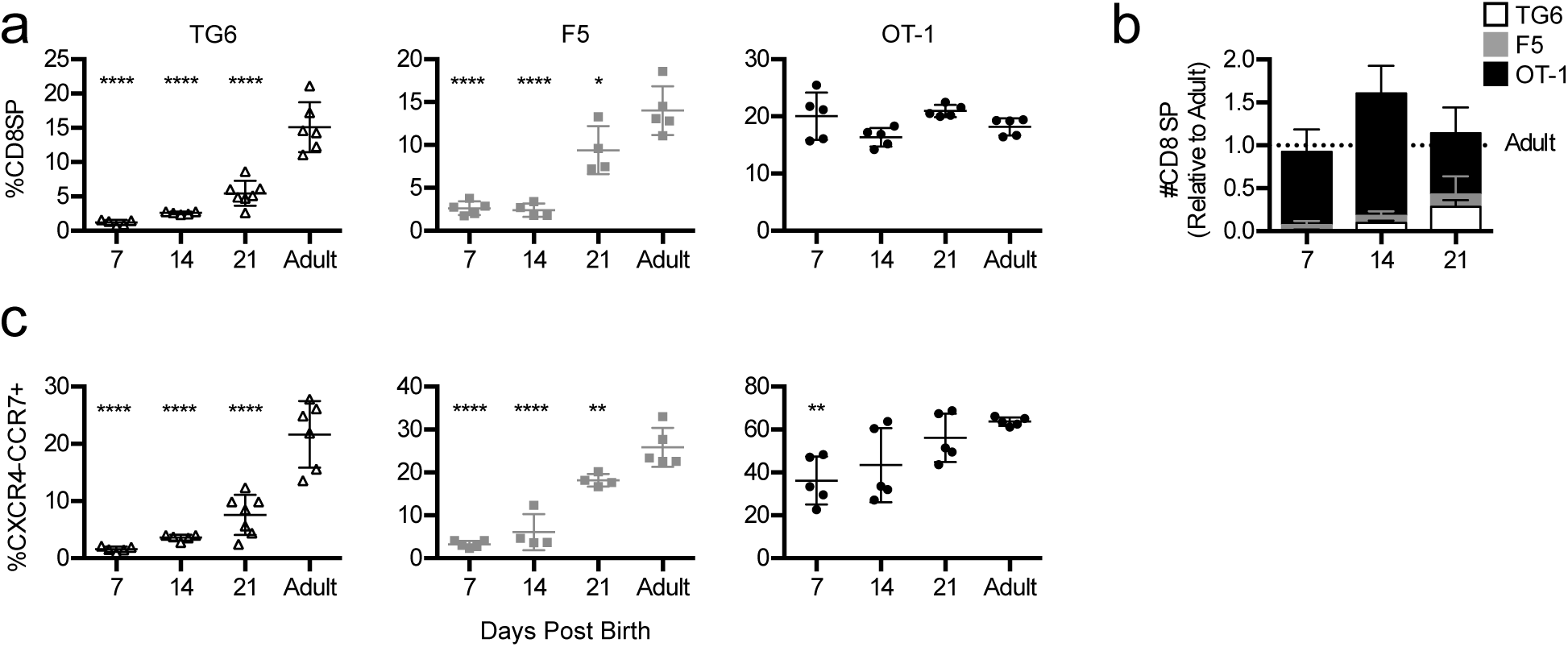
Mature thymocytes with lower self-reactivity appear later post birth than those with higher self-reactivity. Thymuses were harvested from TG6 (triangle, white bar), F5 (square, grey bar), or OT-1 (circle, black bar) transgenic neonatal mice at 7, 14, and 21 days post birth and from adult mice (6-9 weeks old), and analyzed using flow cytometry. (a) Percentage of CD4-CD8+ cells. (b) Total number of CD4-CD8+ thymocytes at the indicated timepoint post birth, relative to adult, for each transgenic. (c) Percentage of CXCR4-CCR7+ cells within CD4+CD8+ and CD4-CD8+ populations. Data are presented as average ± SD and analyzed using an ordinary one-way ANOVA, Dunnett’s multiple comparisons (*P <0.05, **P <0.01, and ****P <0.0001) comparing to adult. All data are compiled from 3 or more experiments.

To examine the timing of positive selection *in vivo* in adult mice, we tracked a cohort of thymocytes using pulse labeling with the thymidine analog 5-ethynyl-2’-deoxyuridine (EdU). *In vivo* EdU injection labels a cohort of pre-positive selection thymocytes proliferating after a successful TCRβ chain rearrangement at the time of the injection, which then can be detected by flow cytometry, as they mature *in vivo* (**Fig. 6a**)(Lucas, Vasseur, and Penit 1993). EdU incorporated equally into the thymocytes of all three TCRtgs (**Fig. 6b**). While the percent of DP cells was greater than the percent of CD8SP cells within the EdU+ population in all three transgenics at 2 days post injection (p.i.) (**Fig. 6c**), EdU+ DP cells in TG6 mice remained higher than EdU+ CD8SP cells until day 6 p.i.. For F5 thymocytes, EdU+ CD8SP overtook the DPs one day earlier (day 5p.i.), and EdU+ CD8SP OT-1 thymocytes accumulated even more quickly—by day 4p.i. (**Fig. 6c**). Though both F5 and OT-1 thymocytes reach a plateau of EdU+ SP by day 5 or 6, TG6 DP thymocytes are still developing into CD8SP thymocytes 7 to 9 days p.i. This data shows that increased time to complete positive selection is also correlated with low self-reactivity *in vivo*. Furthermore, these results show that some CD8SP thymocytes can take longer than 7 days to complete positive selection, as also suggested by earlier studies(Saini et al. 2010; Lucas, Vasseur, and Penit 1993).

**Figure 6:**
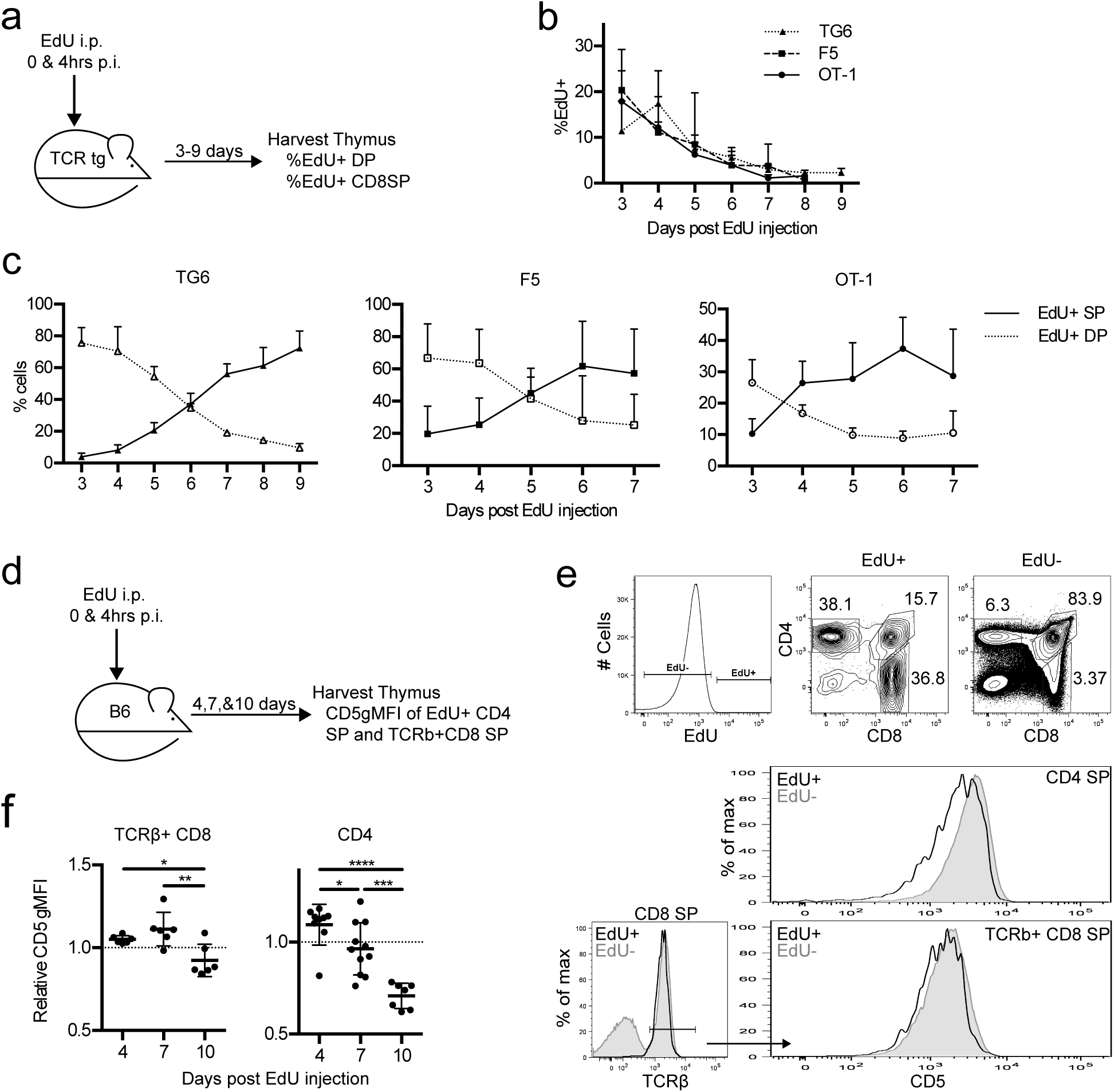
Thymocytes with lower self-reactivity have delayed development in steady state TCRtg and wildtype mice. TG6, F5, or OT-1 (a,b,c) or B6 (d,e,f) mice were injected with two doses of 1mg of EdU intraperitoneally (i.p.) at 0 and 4 hours post injection (p.i.). (a) TCRtg experimental schematic. (b) EdU incorporation into TCRtg thymuses. (c) Percent CD8SP and DP thymocytes out of gated EdU+ cells at various days p.i. in TG6, F5, and OT-1 tg thymuses (left to right). (d) B6 experimental schematic. (e) Representative gating strategy and histograms showing B6 day 10 p.i. CD5 expression on EdU+ and EdU-mature SP cells. (f) CD5 expression of EdU+TCRβ+CD8SP and EdU+ CD4SP thymocytes at 4, 7, and 10 days p.i.. CD5 expression shown relative to total TCRb+CD8SP or CD4SP CD5 expression. For (f) data are presented as average ±SD and analyzed using an ordinary one-way ANOVA, Tukey’s multiple comparisons (** P<0.01, ***P<0.001, ****P<0.0001). All data are compiled from 3 or more experiments.

To confirm the relationship between self-reactivity and time to complete positive selection, we also performed EdU pulse experiments in non-TCRtg mice. We pulsed wild-type mice with EdU and then examined CD5 levels on EdU+ mature TCRβ+CD8 and CD4SP thymocytes at 4, 7, and 10 days p.i. (**Fig. 6d**). We observed that EdU+ mature thymocytes found at earlier time points (and that therefore completed positive selection earlier) have higher CD5 compared to EdU+ mature thymocytes found at later timepoints (**Fig. 6e** and **f**). This is true for TCRβ+CD8SP thymocytes, however the trend is even more dramatic in CD4SPs (**Fig. 6f**). These data indicate that self-reactivity and time to complete positive selection are also inversely correlated in polyclonal mice, and this relationship exists for both MHC-I-restricted and MHC-II-restricted thymocytes.

### Thymocytes with low self-reactivity retain a preselection gene expression pattern marked by elevated expression of ion channel genes

Positive selection is accompanied by large scale changes in gene expression, and delayed progress through positive selection for thymocytes with low self-reactivity might lead to synchronous or asynchronous delays in these gene expression changes. To investigate this question, we performed bulk RNA sequencing (RNA-seq) on thymocytes from TG6, F5, and OT-1 mice after sorting into three stages of development: early positive selection (CD4+CD8+CXCR4+CCR7-), late positive selection (CD4+CD8+CXCR4-CCR7+), and mature CD8SP (CD4-CD8+CXCR4-CCR7+). TG6 and OT-1 thymocytes showed significant differences in gene expression at each developmental stage, with the number of differentially expressed genes decreasing with maturity (**Fig. S7**). Gene set enrichment analysis (GSEA) revealed that gene sets whose enrichment was associated with high self-reactivity (i.e., OT-1>F5>TG6) were related to ribosome function, RNA processing, and translation in early and late positive selection thymocytes (**Fig. 7a**, top, and **S7**, red dots). Genes involved in cell division (**Fig. S7**, purple dots) and effector function (**Fig. S7**, blue dots) were also upregulated in mature CD8SP thymocytes with high self-reactivity (**Fig. 7a**). Intriguingly, gene sets whose enrichment was associated with low self-reactivity (i.e., TG6>F5>OT-1) were related to transmembrane ion channel function—particularly voltage gated ion channels (**Fig. 7b**, and **Fig S7**, green dots). Genes contributing towards the enrichment scores of ion channel related gene sets, or leading-edge (LE) genes, included components of calcium (*Cacna1e, Cacnb1, Cacnb3, Cacng4*), potassium (*Kcna2, Kcna3, Kcnh2, Kcnh3*), sodium (*Scn2b, Scn4b, Scn5a*), and chloride (*Clcn2*) ion channels.

**Figure 7:**
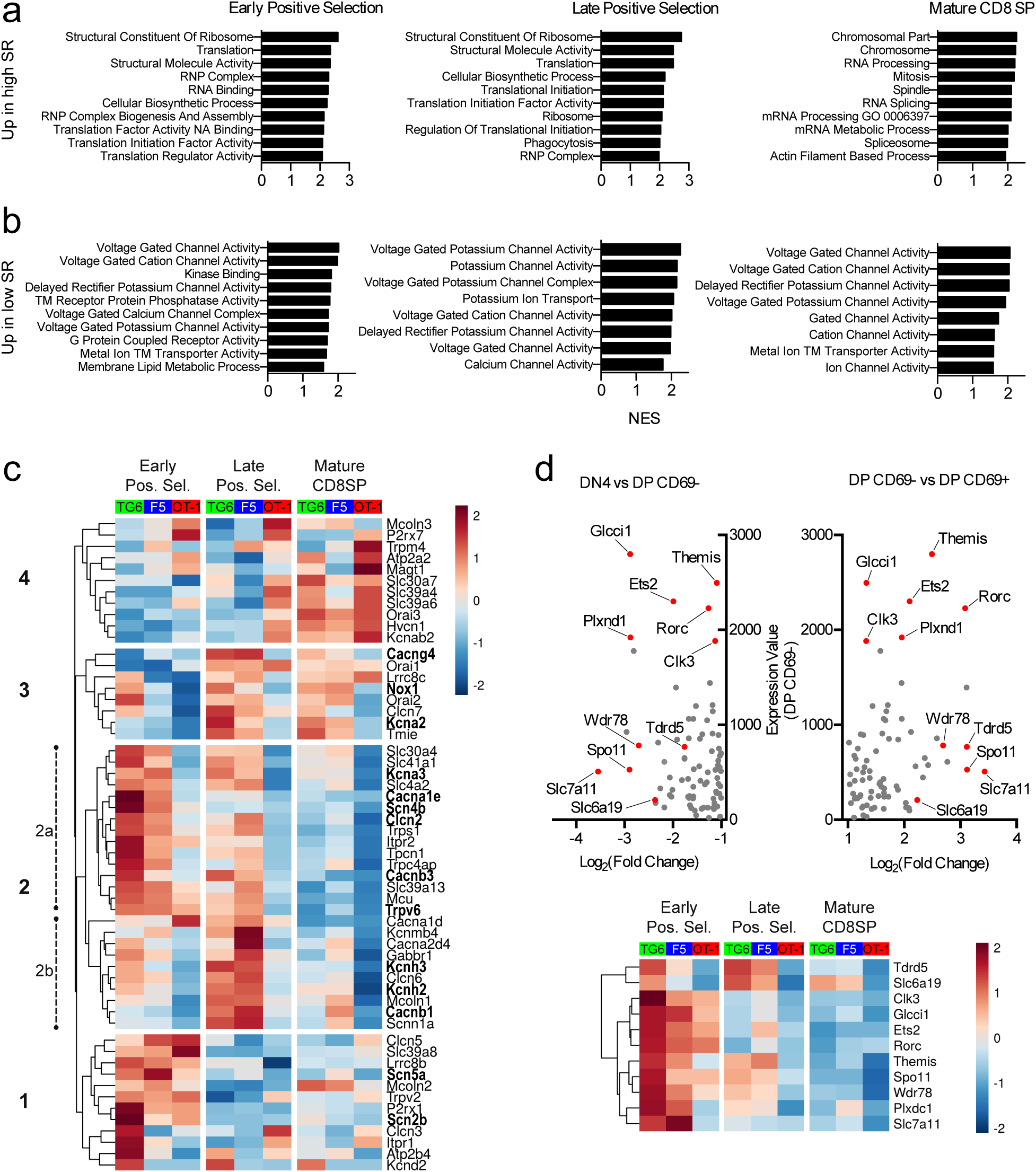
Thymocytes with low self-reactivity retain a preselection gene expression program marked by elevated expression of ion channel genes. RNA-seq of OT-1, F5, and TG6 thymocytes at the early positive selection, late positive selection, and mature CD8SP stages. All samples have three biological replicates except for TG6 late positive selection, which has two biological replicates. (a and b) Gene set enrichment analysis (GSEA) was performed on all pairwise combinations of TCRtg samples and the gene sets were filtered for enrichment in the order of self-reactivity (OT-1 > F5 > TG6) (a) or in the inverse order of self-reactivity (TG6 > F5 > OT-1) (b). Normalized enrichment scores (NES) are from the OT-1 vs TG6 (a) or TG6 vs OT-1 (b) comparison. Abbreviations: self reactive (SR), ribonucleoprotein (RNP), nucleic acid (NA), transmembrane (TM). (c) Heatmap of differentially expressed (padj<0.05 for OT-1 vs TG6 comparison) ion channel genes (see methods) across all three stages and all three transgenic models. Gene expression values normalized by the DESeq2 variance stabilizing transformation were averaged over biological replicates and scaled per row. Heatmap is hierarchically clustered into four groups, numbered in the order of expression during development. Leading edge genes from GSEA are in bold. (d) (Top) Plot of expression level in DP CD69-thymocytes versus fold difference for DN4 versus DP CD69-(left side of X-axis) or DP CD69-versus DP CD69+ (right side of X-axis) for ImmGen microarray data. The eleven genes chosen to represent the “preselection DP” gene signature are indicated with red dots. (Bottom) Heatmap of normalized expression (as in (c)) of the 11 representative preselection DP genes in TG6, F5, and OT-1 thymocytes at the indicated stage of development.

To obtain a broader perspective on the relationship between thymocyte self-reactivity and ion channel gene expression, we curated a list of 56 ion channel genes expressed in thymocytes and examined expression changes associated with thymocyte maturation stage and self-reactivity (**Fig. 7c and S8a**). At all three developmental stages, there were more ion channel genes with higher gene expression in TG6 (green bars) compared to OT-1 (red bars) thymocytes **(S8a)**. Hierarchical clustering based on gene expression patterns across all samples revealed a group of genes (group 2a) that shows a strong association with low self-reactivity, and is also upregulated during early positive selection within each mouse strain. This cluster includes *Scn4b* (**Fig. S8b**), which encodes the regulatory subunit of a VGSC linked to positive selection(Lo, Donermeyer, and Allen 2012). Groups 1, 2b, and 3 are also upregulated in cells with low self-reactivity but their expression peaks later during positive selection. Eleven out of the 56 genes (group 4) have higher expression in cells with high self-reactivity and also peak at the mature CD8SP stage. This group includes *P2rx7* and *Trmp4*, both of which have been linked to modulation of T cell effector responses(Trebak and Kinet 2019; Feske, Wulff, and Skolnik 2015).

The observation that many of the ion channel genes that correlated with low self-reactivity were also preferentially expressed at the early positive selection stage (**Fig. 7c**, group 2a) suggested that thymocytes of low self-reactivity may retain an immature gene expression pattern later into positive selection. To investigate this question, we used microarray data from the Immunological Genome Project (ImmGen)(Heng, Painter, and Consortium 2008) to identify a representative set of eleven genes preferentially expressed in non-TCR transgenic, preselection (CD69-) DP thymocytes relative to postselection (CD69+) DP thymocytes, and to their immediate precursor (DN4 thymocytes) (see methods) (**Fig. 7d**, top). We then examined expression of these genes in TG6, F5, and OT-1 thymocytes at different developmental stages (**Fig. 7d**, bottom). Interestingly, all eleven genes correlated with low self-reactivity, particularly at the early DP stage of development. This included *Themis* (**Fig. S8b**), which encodes a protein required for modulating TCR signaling during positive selection(Lesourne et al. 2009; Patrick et al. 2009; Kakugawa et al. 2009; Fu et al. 2009; Johnson et al. 2009; Choi, Warzecha, et al. 2017; Choi, Cornall, et al. 2017; Mehta et al. 2018). Moreover, expression of the curated set of ion channel genes in the ImmGen microarray data set revealed that group 2 genes, whose expression correlated with low self-reactivity, also displayed higher expression in preselection wildtype thymocytes (**Fig S8c**). Furthermore, GSEA between CD69-DP compared to DN4 or CD69+ DP thymocytes showed enrichment of gene sets relating to ion channels (DN4 < CD69-DP > CD69+ DP) (**Fig. S8d**). Together, this data suggests that cells with low self-reactivity retain a preselection DP phenotype later into development than those with high self-reactivity, and that elevated expression of ion channel genes is a prominent feature of that phenotype.

Retention of a preselection gene expression program can account for some, but not all, of the elevated ion channel gene expression by thymocytes of low self-reactivity. Specifically, there is another set of ion channel genes whose expression peaks at the late positive selection or CD8SP stage (**Fig. 7c**, group 3). Interestingly, two of these genes, *Kcna2* and *Tmie* (**Fig. S8b**), are also upregulated in mature peripheral CD5 low CD8SP T cells relative to CD5 high CD8SP T cells, based on published data sets (**Table S3** and **S4**)(Fulton et al. 2015; Matson et al. 2020; Barrett et al. 2013). Additionally, cells with low self-reactivity also have high expression of the sodium-dependent neutral amino acid transporter *Slc6a19* (**Fig. S7** and **S8b**, **Table S3** and **S4**)(Fulton et al. 2015; Matson et al. 2020). Thus, elevated expression of distinct sets of ion channel genes is observed throughout the development of CD8 T cells with low self-reactivity.

## Discussion

T cell positive selection is a multi-day process during which thymocytes experience transient, serial TCR signals, triggering a series of phenotypic changes culminating in co-receptor down regulation and lineage commitment. Preselection DP thymocytes are highly sensitive to low affinity self-ligands, and gradually downmodulate their sensitivity as they mature(Davey et al. 1998; Lucas et al. 1999). However the factors that modulate TCR sensitivity during positive selection are not well understood. Moreover, there is a growing appreciation that thymocytes differentially adjust their response to self-ligands based on their degree of self-reactivity, although little is known about how positive selection differs for thymocytes of relatively low versus relatively high self-reactivity. Here we show that thymocytes with low self-reactivity experience briefer TCR signals and can take more than twice as long to complete positive selection, compared to those with relatively high self-reactivity. Thymocytes with low self-reactivity retain a preselection gene expression program as they progress through development, including elevated expression of genes previously implicated in modulating TCR signaling during positive selection, and a novel set of ion channel genes. In addition, a separate set of ion channel genes that peaks at the CD8SP stage is also selectively upregulated in mature CD8SP thymocytes of low self-reactivity. These results indicate that the modulation of TCR sensitivity during T cell development occurs with distinct kinetics for T cells with low self-reactivity, and suggests that changes in membrane ion channel activity may be an important component of T cell tuning during both early and late stages of T cell development.

Our results contrast with an earlier study, which reported that the overall time for positive selection was constant for different CD8 T cell clones, with an accelerated early phase compensating for a delayed late phase for CD8 T cells of low self-reactivity(Kimura et al. 2016). Importantly, that study relied on the decrease in Rag-GFP reporter expression to infer the timing of development, rather than directly tracking cohorts of positively selecting cells as we have done in the current study. While the slow loss of Rag-GFP reporter due to GFP protein turnover can provide an indication of the elapsed time after shutoff of rag expression(McCaughtry, Wilken, and Hogquist 2007), reporter expression during the early phase of positive selection may be impacted by the timing of rag shutoff and cell turnover, making this an unreliable indicator of developmental time in this experimental setting.

A variety of molecules have been implicated in modulating the TCR sensitivity of DP thymocytes, including the TCR-associating protein Themis, ER associating protein Tespa1, microRNA mir-181a, and components of a VGSC (encoded by *Scn4b* and *Scn5a*)(Wang et al. 2012; Lo, Donermeyer, and Allen 2012; Fu et al. 2009; Johnson et al. 2009; Lesourne et al. 2009; Patrick et al. 2009; Kakugawa et al. 2009). Here we show that the VGSC is part of a larger group of ion channel genes, that also includes components of voltage gated calcium channels (*Cacna1e, Cacnb3*), that are both upregulated in preselection DP thymocytes, and selectively retained during the positive selection of thymocytes with low self-reactivity. We also identified another group of ion channels whose expression peaks at the CD8SP stage, and that are also upregulated in CD8 T cells of low self-reactivity. This group includes *Kcna2,* a component of a voltage gated potassium channel, and *Tmie* a gene essential for calcium-dependent mechanotransduction in hair cells within the inner ear(Qiu and Müller 2018; Zhao et al. 2014). Thus, ion channel expression appears to be selectively modulated based on the degree of self-reactivity at two distinct stages of T cell development, and may help to enhance weak TCR signals during both the initial phase of positive selection and during T cell homeostasis in the periphery.

Voltage gated ion channels are best known for their role in generating action potentials in neurons and muscle cells, but they also play roles in non-electrically excitable cells such as T cells and thymocytes(Feske, Wulff, and Skolnik 2015; Lo, Donermeyer, and Allen 2012; Cahalan and Chandy 2009; Trebak and Kinet 2019). In these cells, voltage gated ion channels have been proposed to be regulated by local fluctuations of membrane charge, or alternative mechanisms such as phosphorylation(Rook et al. 2012). While the link between ion channels and T cell tuning is not known, the modulation of calcium entry is a potential mechanism. TCR triggering initially leads to a rise in cytosolic calcium via release of ER stores, and the calcium rise is sustained in mature T cells when depletion of ER calcium stores triggers the opening of calcium release activated channels (CRAC). CRAC channels appear to be dispensable for positive selection(Oh-Hora 2009; Gwack et al. 2008; Vig and Kinet 2009), and it remains unclear how thymocytes sustain calcium signals during the days required to complete positive selection. Evidence suggests that the VGSC promotes prolonged calcium flux in response to TCR triggering in DP thymocytes, although the mechanism remains unclear(Lo, Donermeyer, and Allen 2012). Our data indicates that, not only sodium, but also potassium, chloride, and calcium channels are upregulated in preselection DP thymocytes and positive selecting thymocytes with low self-reactivity, implying that positive selection leads to broad changes in transmembrane ion flux. In addition to potentially promoting calcium influx, changes in membrane ion flux could also directly impact TCR triggering by regulating local membrane charge at the receptor complex(Ma et al. 2017). Altered ion flux in T cells with low self-reactivity may also impact ion-coupled transport of nutrients. In this regard, it is intriguing that the sodium-dependent neutral amino acid transporter gene *Slc6a19* is also consistently upregulated in thymocytes and CD8 T cells with low self-reactivity (**Fig. 7c** and **d**, **S7** and **Table S3** and **S4**)(Fulton et al. 2015; Matson et al. 2020).

Taken together, our results support a model in which TCR tuning based on the degree of self-reactivity occurs in two temporally-distinct phases (**Fig. 8**). Thymocytes with low self-reactivity (**Fig. 8**, dashed lines) experience briefer TCR signals and accumulate TCR signaling more slowly, resulting in a delayed time course of positive selection. During phase 1, a “preselection DP” gene expression program (including *Themis, Scn4b*, and *Cacna1e*) is down-regulated in all thymocytes undergoing positive selection, but more gradually and incompletely in thymocytes with low self-reactivity. During phase 2, a distinct set of genes (including *Kcna2* and *Tmie*) whose expression peaks in CD8 SP, are upregulated to a greater extent in thymocytes with low self-reactivity. Higher expression of both sets of genes help to sustain TCR signals throughout positive selection, allowing thymocytes to continue to receive survival and differentiation signals in spite of their low self-reactivity.

**Figure 8:**
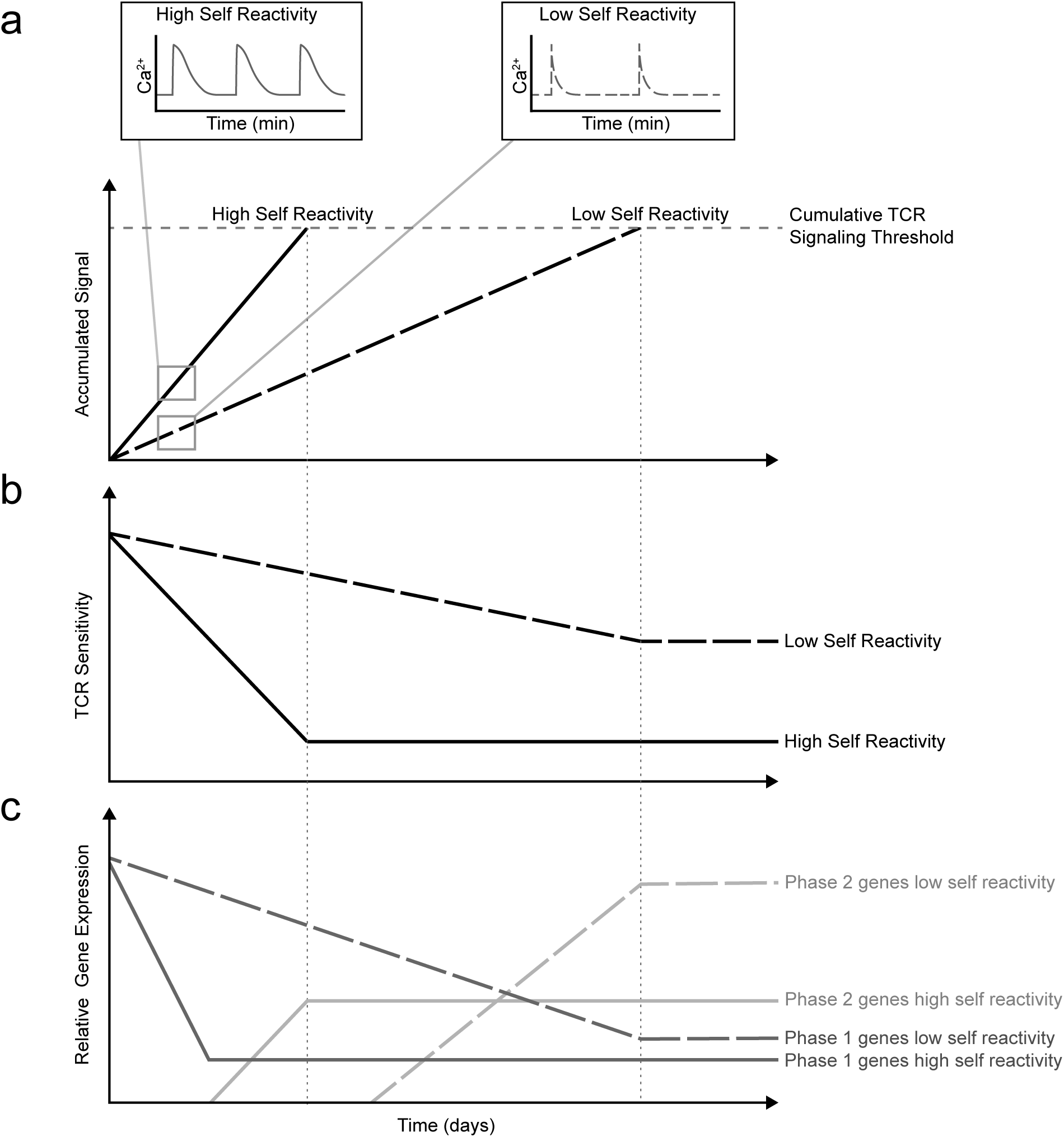
A model for temporal signal accumulation and TCR sensitivity during development in cells with relatively high and low self-reactivity. (a) Positively selected cells with relatively high self-reactivity (SR) receive serial, brief (minutes) TCR signals, allowing for signal accumulation over a period of days, to eventually complete positive selection. Those with relatively low self-reactivity (dashed lines) experience briefer TCR signals leading to slower signal accumulation and a longer time to achieve the cumulative TCR signaling threshold required for positive selection. (b) All DP thymocytes begin positive selection with high TCR sensitivity, and gradually downregulate their sensitivity as they accumulate TCR signals. Thymocytes with low self-reactivity downregulate their sensitivity more gradually and less completely than those with high self-reactivity. (c) An early (phase 1: black lines) and a late (phase 2: grey lines) set of gene expression changes modulate TCR signaling during positive selection. Elevated expression of phase 1 genes (including *Themis, Scn4b*, and *Cacna1e*) help to confer high TCR sensitivity on preselection DP thymocytes. Phase 1 genes are downregulated more slowly in thymocytes with low self-reactivity, allowing them to retain high TCR sensitivity later into development, and aiding in signal accumulation from weak ligands. Later during positive selection, phase 2 genes (including *Kcna2* and *Tmie*) are upregulated to a greater extent in thymocytes with low self-reactivity.

Previous work has shown that CD8 T cells can take 3 or more days to complete development, but it was unclear why positive selection takes longer for some cells than others(Saini et al. 2010; Lucas, Vasseur, and Penit 1993; Kurd and Robey 2016). Our data show that positive selection can take considerably longer for cells with low self-reactivity compared to those with high self-reactivity. In particular, EdU pulse labeling in adult TG6 transgenic mice shows that some thymocytes are still converting from DP to CD8SP after more than 1 week of positive selection. It has been previously noted that CD8 T cells that arise in the neonatal period have higher CD5 levels, and are more proliferative and responsive to cytokines compared to CD8 T cells that arise in adult mice(Dong et al. 2017; Smith et al. 2018). This phenomenon may be explained in part by the shorter time required for T cells with high self-reactivity to complete positive selection. However, distinct mechanisms independent of TCR specificity also contribute to this phenomenon. For example, T cells that arise in the neonatal period are derived from fetal stem cells, and are intrinsically programmed to be more proliferative than those derived from adult stem cells(Smith et al. 2018; Wang et al. 2016; Mold et al. 2010; Ikuta et al. 1990; Bronevetsky, Burt, and McCune 2016). There are also indications that age-related changes in the thymic environment may alter the TCR affinity thresholds for positive and negative selection(Dong et al. 2017). Thus, three distinct mechanisms may all contribute to a bias toward high self-reactivity T cells in young individuals, with T cells of lower self-reactivity accumulating with age. It is tempting to speculate that this arrangement is evolutionarily advantageous, since highly self-reactive T cells can provide a rapid response against acute infections, which are the biggest threat to a young individual. As the organism ages, however, it may become advantageous to have a more diverse, specialized T cell repertoire, particularly to fight chronic infections. In line with this notion, we recently reported that TG6 T cells are resistant to exhaustion and provide a protective response to *T. gondii* chronic infection, properties that are linked to limited binding of self-peptides to the restricting MHC-1 molecule L^d^ (Blanchard et al. 2008; Chu et al. 2016; Tsitsiklis et al. 2020).

Overall, the work presented here provides insight on the differences in the development of T cells with low and high self-reactivity that may impact their distinct peripheral functions. The question of how a lengthy positive selection process and differences in ion channel expression help to functionally tune T cells with low self-reactivity remains an important area for future investigations.

## Materials and Methods

### Mice and bone marrow chimeras

All mice were bred and maintained under pathogen-free conditions in an American Association of Laboratory Animal Care-approved facility at the University of California, Berkeley. The University of California, Berkeley Animal Use and Care Committee approved all procedures. B6 (C57BL/6, Stock No.: 000664), B6.C (B6.C-H2^d^/bByJ, Stock No.: 000359), and Rag2^−/−^ (B6(Cg)-Rag2^tm1.1Cgn^/J, Stock No.: 008449), Rag1^−/−^ (B6.129S7-Rag1^tm1Mom^, Stock No.: 002216), and B2M^−/−^ (B6.129P2-B2m^tm1Unc^/DcrJ, Stock No.: 002087) mice were from Jackson Labs. OT-1 Rag2^−/−^ (B6.129S6-Rag2^tm1Fwa^ Tg(TcraTcrb)1100Mjb, Model No.: 2334) mice were from Taconic. F5 Rag1^−/−^ and B6xB6C (H2^b/d^) mice were generated by crossing as previously described(Au-Yeung et al. 2014; Chu et al. 2016). TG6 transgenic mice were generated as previously described by Chu et al., 2016(Chu et al. 2016), and further crossed in house to generate TG6 H2^b/d^ (used in neonatal, EdU pulse experiments, and RNA-seq), and TG6 H2^b/d^ Rag2^−/−^ (used to measure self-reactivity), and TG6 H2^b^ Rag2^−/−^ mice (used for overlay onto thymic slices). Preselection OT-1 Rag2^−/−^ thymocytes were generated by crossing onto a non-selecting MHC-I deficient background (B2M^−/−^), or by transferring 1 × 10^6^ OT-1 Rag2^−/−^ bone marrow cells intravenously (i.v.) into lethally irradiated (1,200 rad) β2M^−/−^ recipients (used for imaging experiments). Preselection F5 Rag1^−/−^ thymocytes were generated by transferring 1 × 10^6^ bone marrow cells i.v. into lethally irradiated (1,200 rad) β2M^−/−^ recipients. Preselection TG6 Rag2^−/−^ thymocytes were generated by crossing onto a non-selecting background lacking H2^d^ MHC-I (B6, H2^b^ haplotype). Bone marrow chimeras were analyzed 5–7 weeks following reconstitution. For all experiments, mice were used from three to ten weeks of age, with the exception of neonatal experiments where mice were used one to nine weeks of age.

### Experimental Design

Sample size in this study was consistent with previous studies, and was not determined based on a prior statistical test. All experiments were completed with at least 3 biological replicates unless otherwise stated in the figure legend. Consistent with previous studies, 3-8 thymic slices (technical replicates) per condition were used for thymic slice experiments analyzed by flow cytometry. Samples were excluded if their thymic phenotype indicated signs of stress (<10^7^ total cells or less than 20% DP thymocytes out of live thymocytes). Samples for thymic slice experiments were excluded if the culture had very low viability (<15%) or <0.2% of donor derived thymocytes of total live cells in the final sample. For EdU experiments, samples that did not incorporate detectable EdU (<0.001% of live cells that were EdU+) were excluded. For experiments where thymic slices from the same genotypic background received multiple treatments, slices were randomly allocated to treatment conditions.

### Thymic slices

Preparation of thymic slices has been previously described(Dzhagalov et al. 2012; Ross et al. 2015). Briefly, thymic lobes were gently isolated and removed of connective tissue, embedded in 4% agarose with a low melting point (GTG-NuSieve Agarose, Lonza), and sectioned into slices of 400-500μm using a vibratome (VT1000S, Leica). Slices were overlaid onto 0.4μm transwell inserts (Corning, Cat. No.: 353090) set in 6 well tissue culture plates with 1ml cRPMI under the insert. 2×10^6^ thymocytes in 10μl cRPMI were overlaid onto each slice and allowed to migrate into the slice for 2-3 hours. Afterwards, excess thymocytes were removed by gentle washing with PBS and cultured at 37°C 5%CO_2_ until harvested for flow cytometry or two-photon analysis. For flow cytometry, thymic slices were dissociated to single-cell suspensions and then stained with fluorophore-conjugated antibodies. For two-photon imaging, thymic slices were glued onto a glass coverslip and imaged as described below.

### Thymocyte labeling for overlay onto thymic slices

Thymuses collected from TCRtg preselection mice and dissociated through a 70μm cell strainer to yield a cell suspension. For overlay onto thymic slices followed by flow cytometry, TCRtg preselection thymocytes were labeled with 1μM Cell Proliferation Dye eFluor450 or 0.5μM Cell Proliferation Dye eFluor670 as previously described(Kurd et al. 2019). For two-photon imaging of thymic slices, thymocytes were labeled with 2 mM leakage-resistant Indo-1 (Thermo Fisher Scientific, Cat. No.: I1226) as previously described followed by overlay onto slices(Melichar et al. 2013).

### Flow cytometry

Thymic slices, whole thymuses, and spleens were dissociated into FACS buffer (0.5% BSA in PBS) or RPMI and passed through a 70μm filter before staining. Splenocytes were then red blood cell lysed using ACK lysis buffer (0.15M NH_4_Cl, 1mM KHCO_3_, 0.1mM Na_2_EDTA) for 5 minutes at room temperature prior to staining. For thymic slices, cells were first stained for 60 minutes in a 37°C water bath with CXCR4 (L276F12 or 2B11) and CCR7 (4B12) antibodies in 2.4G2 supernatant. Cells were then surfaced stained for 15 minutes on ice in 2.4G2 supernatant containing the following antibodies: CD4 (clone GK1.5 or RM4-5), CD8α (53-6.7), CD5 (53-7.3), and CD69 (H1.2F3). Finally, cells were washed in PBS and stained in Ghost Dye Violet 510 (Tonbo Biosciences, Cat. No.: 13-0870-T100) for 10 minutes on ice. The following antibodies were used for flow cytometric analysis of whole thymuses and spleens: CD4 (GK1.5 or RM4-5), CD8α (53-6.7), CD5 (53-7.3), and CD69 (H1.2F3), CD24 (M1/69), TCRb (H57-597), CXCR4 (L276F12 or 2B11), CCR7 (4B12), Vα2 (B20.1), Vβ8.1/8.2 (KJ16-133.18), and Vβ2 (B20.6). Cells were then washed in PBS and stained in Ghost Dye Violet 510 as described above. For intracellular staining, cells were then fixed and permeabilized using Invitrogen’s Transcription Factor Staining Buffer Set (Thermo Fisher Scientific; Cat. No.: 00-5523-00) according to manufacturer’s instructions and stained with Nur77 (12.14) antibody (Thermo Fisher Scientific, Cat. No.: 12-5965-82). All antibodies were from Thermo Fisher Scientific, Biolegend, or Tonbo Biosciences. Cells were analyzed using an LSRII, LSR, or Fortessa X20 flow cytometer (BD Biosciences) and data analyzed using FlowJo software (Tree Star).

### EdU pulse labeling

TCRtg and non-transgenic selecting mice, aged 5-8 weeks, received two intraperitoneal (i.p.) injections of 5-ethynyl-2’-deoxyuridine (EdU) (Thermo Fisher Scientific, Cat. No.: A10044) 1mg each and 4 hours apart, as previously described for BrdU(Lucas, Vasseur, and Penit 1993). Thymuses were harvested from pulsed mice 3-10 days after the second injection and dissociated into a single-cell suspension and were surface stained with CD4, CD8α, CD5, CD69, CD24, and TCRβ antibodies as described above. Cells were then processed using Click-iT™ EdU Alexa Fluor™ 488 Flow Cytometry Assay Kit from Invitrogen according to the manufacturer’s instructions (Thermo Fisher Scientific, Cat. No.: C10420). Flow cytometry and data analysis was performed as described above.

### Two-photon imaging

Two-photon imaging of thymic slices has been previously described(Melichar et al. 2013; Dzhagalov et al. 2012; Ross et al. 2014; Dzhagalov et al. 2013; Au-Yeung et al. 2014). All imaging was performed 3 to 6 hours after the addition of thymocytes to thymic slices. Thymic slices were glued onto glass cover slips and fixed to the bottom of a petri dish being perfused at a rate of 1mL/min with 37°C oxygenated, phenol red–free DMEM during imaging using an MPII Mini-Peristaltic Pump (Harvard Apparatus) and a 35mm Quick Exchange Platform (QE-1) in conjunction with a TC-324B temperature controller (Warner Instruments). Samples were imaged in the cortex of the thymus; determined by proximity to the thymic capsule. Images were collected using a Zeiss LSM 7 MP upright, two-photon microscope with a 20x/1.0 immersive Zeiss objective and a Coherent Chameleon laser tuned to 720 nm for imaging Indo-1. Ratiometric Ca^2+^ signals were collected with 440nm long pass dichroic mirror and 400/45 and 480/50 bandpass filters. Ca^2+^ hi signals (∼401nm) were collected using a bialkali PMT; Ca^2+^ lo signals (∼475nm) were collected using a GaASP PMT. Image areas of up to 212 x 212 μm to a depth of up to 55 μm were acquired every 10 sec for 15 or 20 min with 3-μm z steps starting from beneath the cut surface, using Zen 2010 software from Zeiss.

### Image analysis

Two-photon movies were processed and rendered using Imaris software (Bitplane Scientific Software) to collect x, y, and z coordinates and fluorescence intensity as previously described(Melichar et al. 2013; Ross et al. 2014; Au-Yeung et al. 2014). Imaging data were then analyzed using a custom MATLAB script (MathWorks; as previously described (Melichar et al. 2013; Ross et al. 2014)), ImageJ, and Microsoft Excel. GraphPad Prism software was used for graphing and statistical analysis. For calculation of the percent of signaling timepoints, data were converted into flow cytometry-like files using custom DISCit software and further analyzed using FlowJo software (TreeStar)(Moreau et al. 2012). The background calcium ratio is average Ca^2+^ ratio under non-selecting conditions (0.675 for OT-1 and TG6) from the Ca^2+^ ratio of the raw fluorescence values at each timepoint per run. To calculate the average event length, cells were considered to be signaling when the average Ca^2+^ ratio was ≥0.2 above the background calcium ratio for ≥2 consecutive timepoints on a cell track. A trigger event was defined by the first of the consecutive timepoints to have a Ca^2+^ ratio ≥0.2 above background. The end of a signaling event was defined by when the Ca^2+^ ratio returned to <0.1 above background for two consecutive timepoints. Only events that had a defined beginning and end were included in calculating signal duration. For tracks with multiple events, only the first signaling event was used to calculating signal duration. For calculating speed over time, we averaged speed values from Imaris over two timepoints (20sec). The speed of signaling and nonsignaling timepoints was measured by averaging the instantaneous speed for each timepoint. The frequency of events was calculated by dividing the number of events compiled from all runs by the cumulative track imaging time (the sum of all of the track durations for all compiled runs), for each condition. For this calculation, events were defined as ≥2 consecutive timepoints of high Ca^2+^ ratio bound by periods ≥4min of nonsignaling.

### RNA preparation, sequencing and analysis

Thymocytes from OT-1 Rag2KO, F5 Rag1KO, and TG6 H2b/d mice were sorted using a BD Influx or BD FACSAria Fusion into CD4+CD8+CXCR4+CCR7-(early positive selection), CD4+CD8+CXCR4-CCR7+ (late positive selection), and CD4-CD8+CXCR4-CCR7+ (Mature CD8SP) populations for RNA sequencing. RNA was harvested using a Quick-RNA Microprep Kit from Zymo Research (Cat. No. R1050) according to the manufacturer’s instructions. RNA integrity was confirmed via Bioanalyzer and Qbit. RNA was sent to BGI Genomics for library generation and RNA sequencing. RNA was sequenced on an Illumina-HiSeq2500/4000 to a depth of 20 million reads.

Sequencing reads were processed with Trimmomatic(Bolger, Lohse, and Usadel 2014) to remove adapter sequences and trim low-quality bases. Reads were aligned to the mm10 genome using Bowtie 2(Langmead and Salzberg 2012) and transcripts were quantified using RSEM(Li and Dewey 2011). Differential expression testing (OT-1 vs TG6, OT-1 vs F5, and F5 vs TG6 at each stage of development) was performed using DESeq2(Love, Huber, and Anders 2014), which produced adjusted p-values corrected by the Benjamini-Hochberg procedure. Gene set enrichment analysis (GSEA) was performed for each DESeq2 comparison using FGSEA(Korotkevich and Sukhov 2019) with gene sets downloaded from MSigDB(Liberzon et al. 2011), including the C5 collection (Gene Ontology (GO) gene sets). Following GSEA, we identified gene sets that were enriched in the order of self-reactivity (OT-1>F5>TG6 or TG6>F5>OT-1) by normalized enrichment score (NES). Normalized counts plotted in Figure S8b were derived by DESeq2’s median of ratios method(Love, Huber, and Anders 2014).

The curated list of ion channel genes was created by first identifying ion channels previously known to be expressed in T cells (207 genes)(Trebak and Kinet 2019; Feske, Wulff, and Skolnik 2015). We narrowed this list to include only genes that were well expressed in our RNA-seq data set (>20 reads in at least one sample), and also significantly different by DEseq2 (padj<0.05) for OT-1 vs TG6 in at least one stage of development. We also included *Tmie*, an ion channel gene differentially expressed by cells with low self-reactivity. This resulted in a list of 56 genes.

### ImmGen microarray analysis

Microarray data (RMA normalized signal intensities) were downloaded from the NCBI GEO database(Edgar, Domrachev, and Lash 2002) from accession number GSE15907(Heng, Painter, and Consortium 2008) and loaded with the GEOquery package(Davis and Meltzer 2007). Samples were filtered to include T.DN4.Th, T.DP.Th, and T.DP69+.Th (three replicates each). Differential expression testing between cell types was performed using limma(Ritchie et al. 2015). For the analysis in Figure 7d, genes representative of a preselection gene expression program were identified from ImmGen microarray data based on differential expression between preselection CD69-DP compared to DN4 and CD69+ DP (log_2_ fold change >1), and either combined expression of >2000 (arbitrary units) (Clk3, Ets2, Glcci1, Rorc, Plxd1, Spo11, Themis) or log_2_ fold change of >2 (Slc6a19, Slc7a11, Tdrd5, Wdr78). Rag1 and Rag2 were excluded. ImmGen populations were relabeled in Figure S8c: T.DN4.Th (DN4), T.DP.Th (DP CD69-), T.DP69+.Th (DP CD69+), T.4SP24-.Th (Mature CD4SP (T)), T.8SP24-.Th (Mature CD8SP (T)), T.4Nve.Sp (Naïve CD4SP (S)), and T.8Nve.Sp (Naïve CD8SP (S)). The gene set enrichment analysis in Figure S8d was performed using FGSEA(Korotkevich and Sukhov 2019) with gene sets downloaded from MSigDB(Liberzon et al. 2011), including the C5 collection (Gene Ontology (GO) gene sets).

### Matson et al. RNA-seq analysis

RNA-seq count data were downloaded from the NCBI GEO database(Edgar, Domrachev, and Lash 2002) from accession number GSE151395(Matson et al. 2020). The data contained four biological replicates for each of four cell types (CD4+CD5lo, CD4+CD5hi, CD8+CD5lo, CD8+CD5hi). Differential expression testing was performed using DESeq2(Love, Huber, and Anders 2014).

### Heatmap Plotting

Heatmaps (**Fig. 7b**, **c**, and **S8c**) were plotted using pheatmap with hierarchical clustering of rows using default settings. Data was scaled per row. For **Figure 7b**, rows (genes) were divided into four clusters based on hierarchical clustering. Genes in **Figure S8c** are arranged by the same ordering as in **Figure 7b**.

### Data Availability

RNA-seq data have been deposited in NCBI’s Gene Expression Omnibus(Edgar, Domrachev, and Lash 2002) and are accessible through GEO Series accession number GSE164896 (https://www.ncbi.nlm.nih.gov/geo/query/acc.cgi?acc=GSE164896).

### Statistical Analysis

Statistical analysis was performed using Prism software (GraphPad) or as explained above for the RNA-seq dataset. Specific statistical tests are indicated in the figure legends. Sample sizes were chosen based on previous publications of similar studies. P values of <0.05 were considered significant.

## Acknowledgements

We would like to thank all the members of the Robey lab for insightful conversation and scientific advice. We would also like to thank Diana Bautista, Nadia Kurd, and Derek Bangs for critical reading of the manuscript. 2-photon imaging experiments were conducted at the CRL Molecular Imaging Center, and we would like to thank Paul Herzmark, Holly Aaron, and Feather Ives for their microscopy training and assistance. This work benefitted from data assembled by the ImmGen consortium. This work was funded by NIH RO1AI064227 (E.A.R.). L.L.M. and A.R.H were supported by NIH T32AI100829, and S.A. was supported by a Human Frontiers Fellowship. Z.S. is supported by the National Science Foundation Graduate Research Fellowship.

## Author Contributions

L.K.L., L.L.M., and E.A.R. designed research; L.K.L., L.L.M., S.P., A.R.H., and S.A. performed research; L.K.L., Z.S., L.L.M., J.K. and E.A.R. analyzed data; and L.K.L. and E.A.R. wrote the paper.

## Supplemental Figures

**Supplemental Figure 1:**
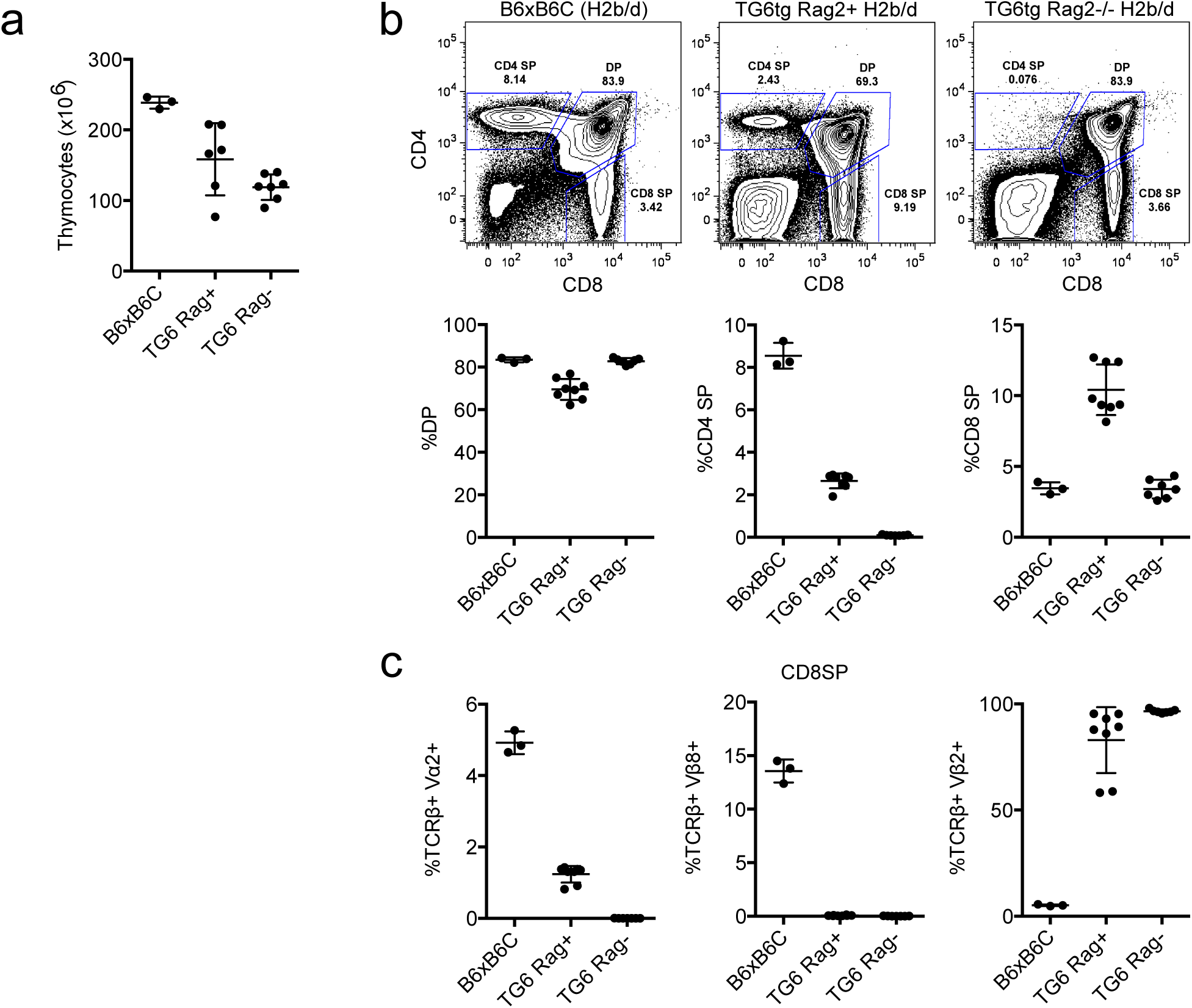
TG6 mice have efficient positive selection and allelic exclusion in the thymus. Thymuses from B6xB6C (H2b/d), TG6 H2b/d Rag+/+, and TG6 H2b/d Rag−/− mice were harvested and analyzed via flow cytometry. (a) Total thymocyte numbers. (b) Representative flow plots and quantification of the percent of DP, CD4SP, and CD8SP thymocytes for the indicated genotypes. (c) Allelic exclusion measured by percent TCRβ and Vα2, Vβ8, or Vβ2 double positive out of CD8SP thymocytes from the indicated genotypes. Data are compiled from two experiments.

**Supplemental Fig. 2:**
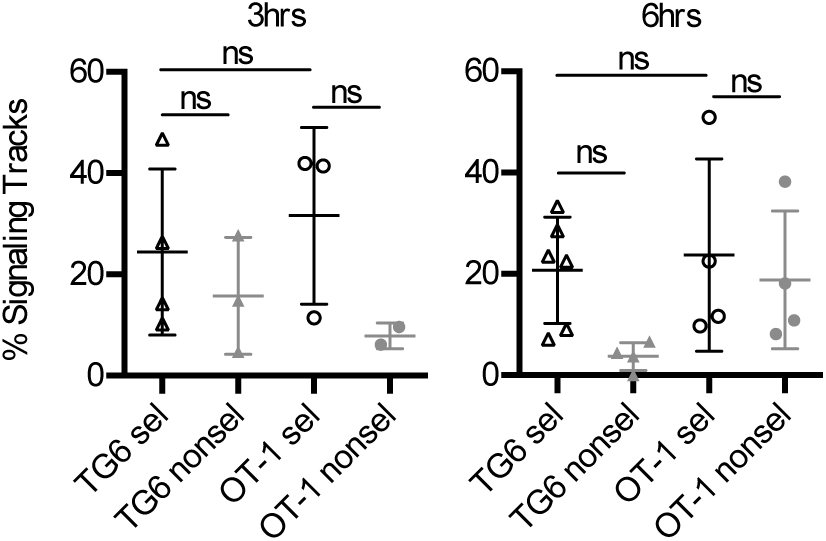
TG6 and OT1 thymocytes have a similar proportion of signaling cells per run. Percentage of signaling tracks (i.e. tracks with at least one signaling timepoint) out of total tracks for each run. Each dot represents a single run. Data are presented as average ± SD and analyzed using an ordinary one-way ANOVA with Tukey’s multiple comparisons (ns=not significant).

**Supplemental Figure 3:**
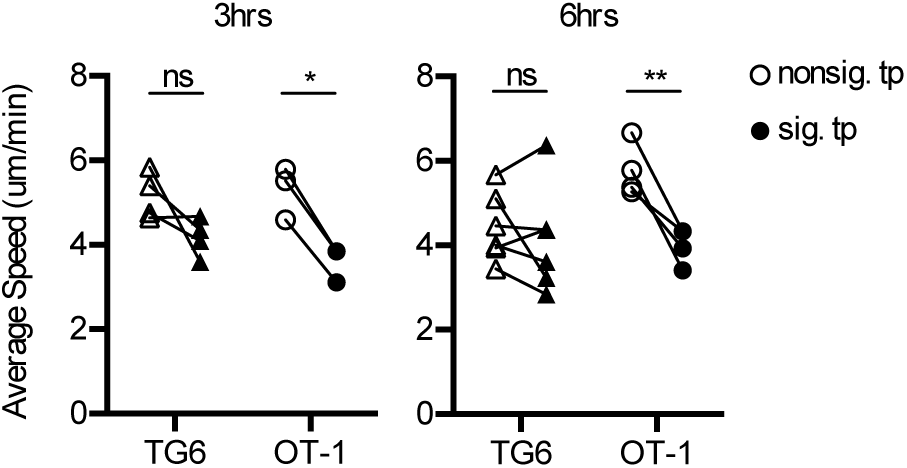
TG6 thymocytes have a less pronounced pause while TCR signaling compared to OT-1 thymocytes. Average instantaneous speed of signaling (sig.) and nonsignaling (nonsig.) timepoints for runs with at least one signaling track. In each column, a dot represents a single run. Between columns, lines connect data from the same run. All data are complied from 2 or more experiments, except OT-1 3hrs data, which is from one experiment. Data are presented as average ± SD and analyzed using an ordinary two-way ANOVA with Sidak’s multiple comparisons (* P<0.05, **P<0.01).

**Supplemental Figure 4:**
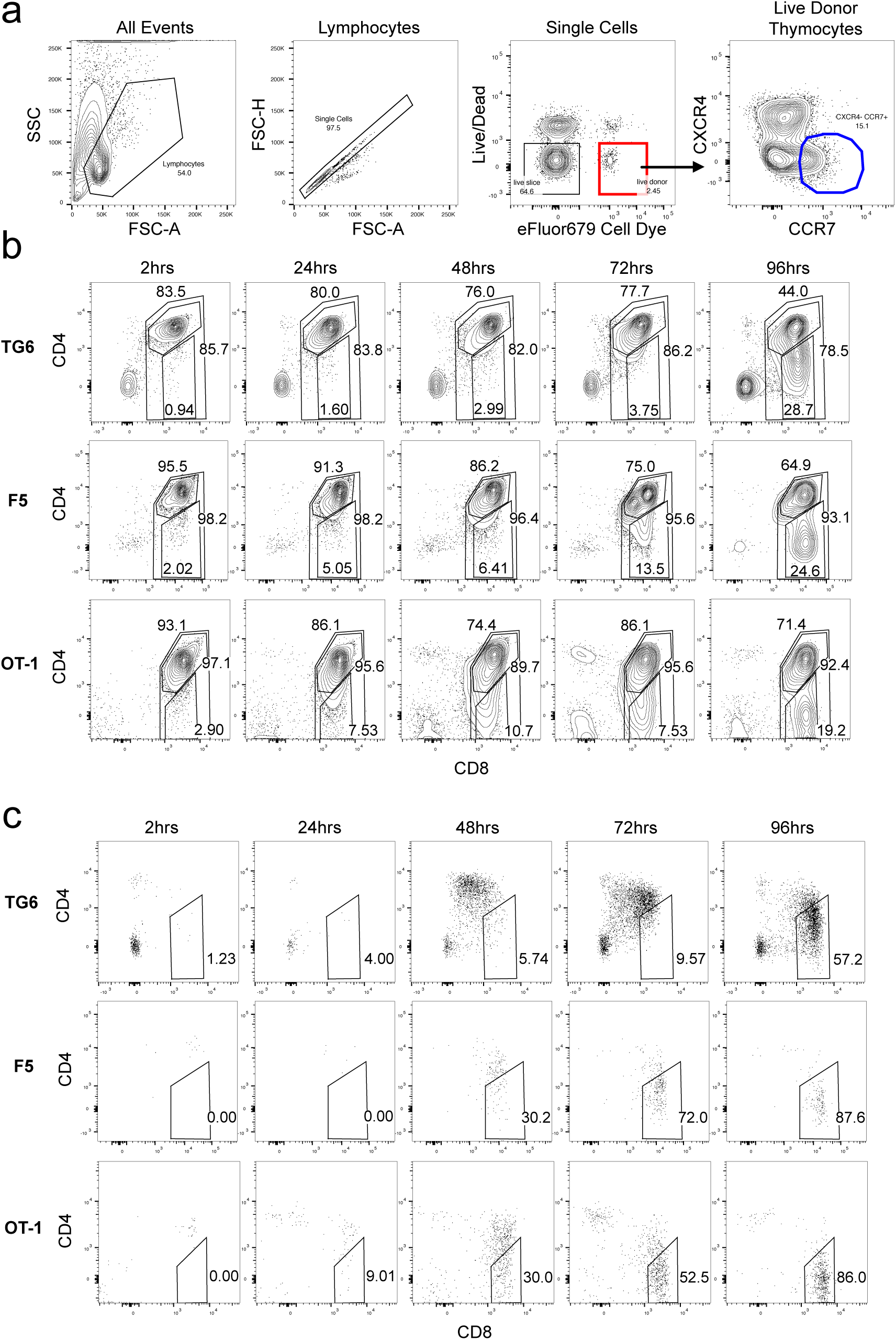
TG6 thymocytes exhibit delayed development in thymic slices. (a) Representative flow plots illustrating the gating strategy for thymic slice experiments. (b) Representative CD4 vs CD8 flow plots of TG6, F5, and OT-1 live donor thymocytes (bolded red gate in a). (c) CD4 vs CD8 representative flow plots for TG6, F5, and OT-1 live donor CXCR4-CCR7+ thymocytes (bolded blue gate in a). Bolded numbers within plots indicate the percentage of cells within a given gate.

**Supplemental Figure 5:**
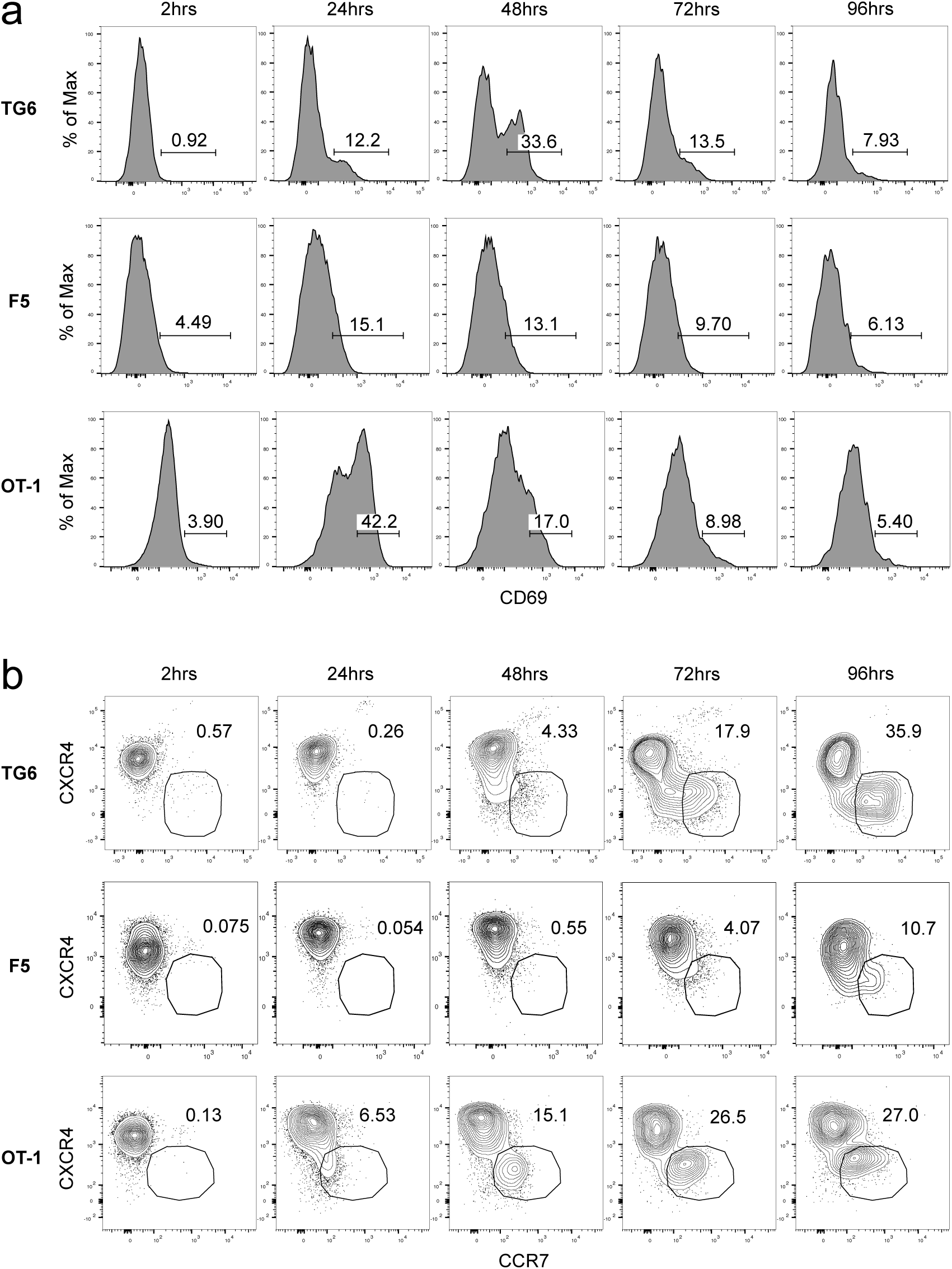
TG6 thymocytes have delayed expression of positive selection markers in thymic slices. Representative flow plots for TG6, F5, and OT-1 thymocytes developing in slices at the indicated timepoints post addition to the slice (timepoint indicated above each column) for (a) CD69 expression and (b) CXCR4 and CCR7 expression. Numbers within plots indicate the percentage of cells within a given gate.

**Supplemental Figure 6:**
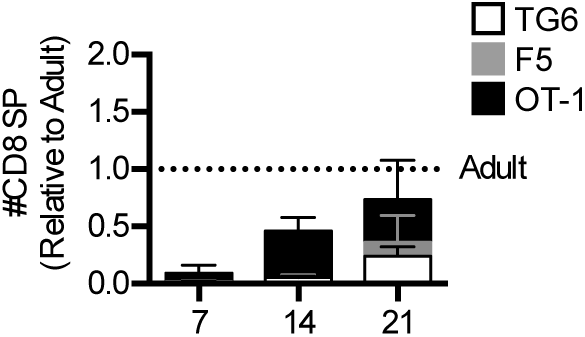
T cells with lower self-reactivity appear later post birth than those with higher self-reactivity in the spleen. Total number of CD4-CD8+ splenocytes relative to adult for each transgenic. Data compiled from 3 or more experiments.

**Supplemental Figure 7:**
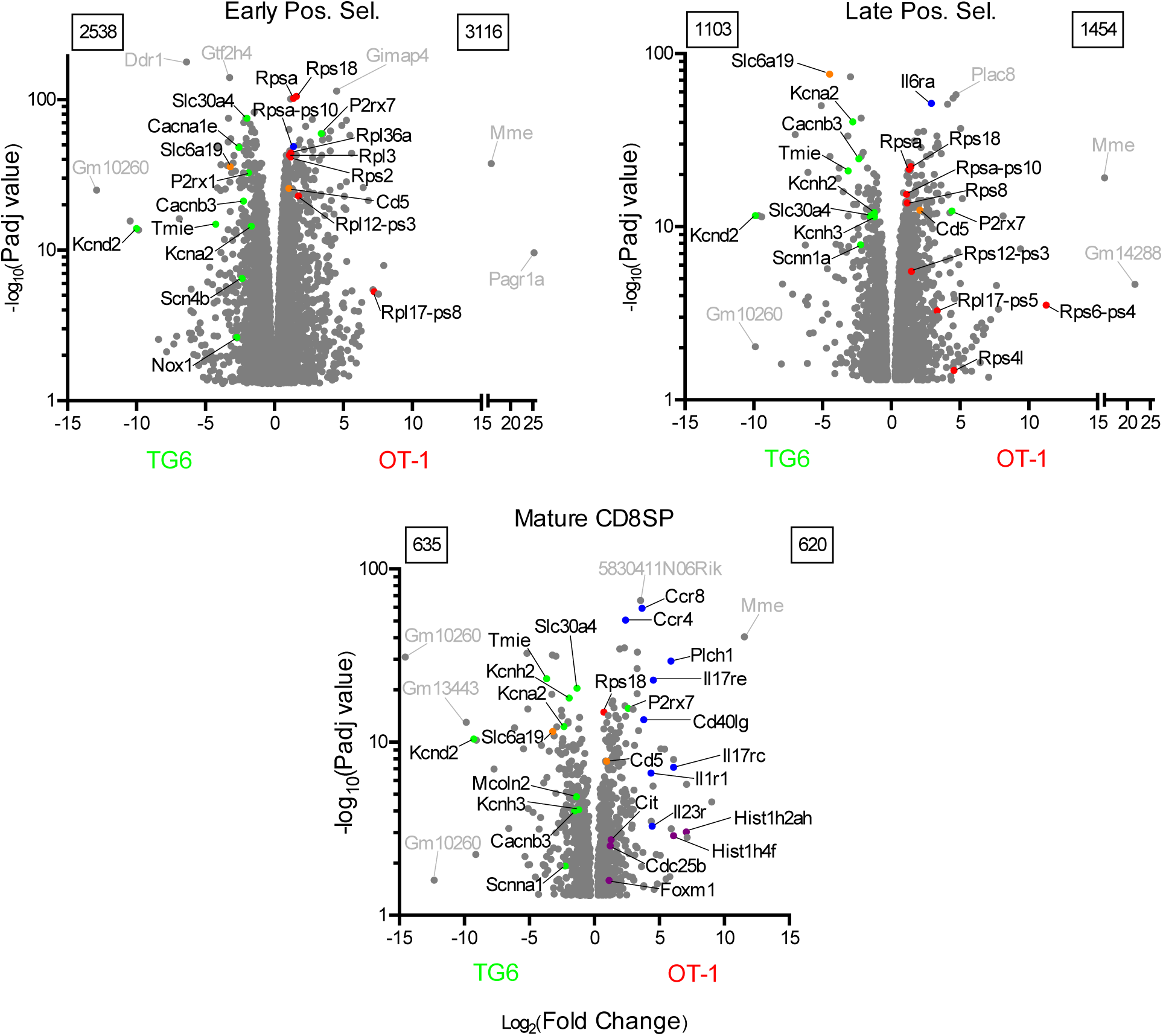
Gene expression differences between TG6 and OT-1 thymocytes at different stages of development. Plots of significance versus fold change for the OT-1 vs TG6 comparison at the early positive selection, late positive selection, and mature SP stages of development. The numbers in boxes denote the number of genes significantly upregulated for TG6 (upper left) and for OT-1 (upper right). Genes discussed in this study are indicated in color: green (ion channel genes), red (translation & ribosome genes), blue (T cell activation/effector function genes), purple (cell division genes), and orange (other). Other genes with very significant p values and/or high fold change are labeled in grey.

**Supplemental Fig. 8:**
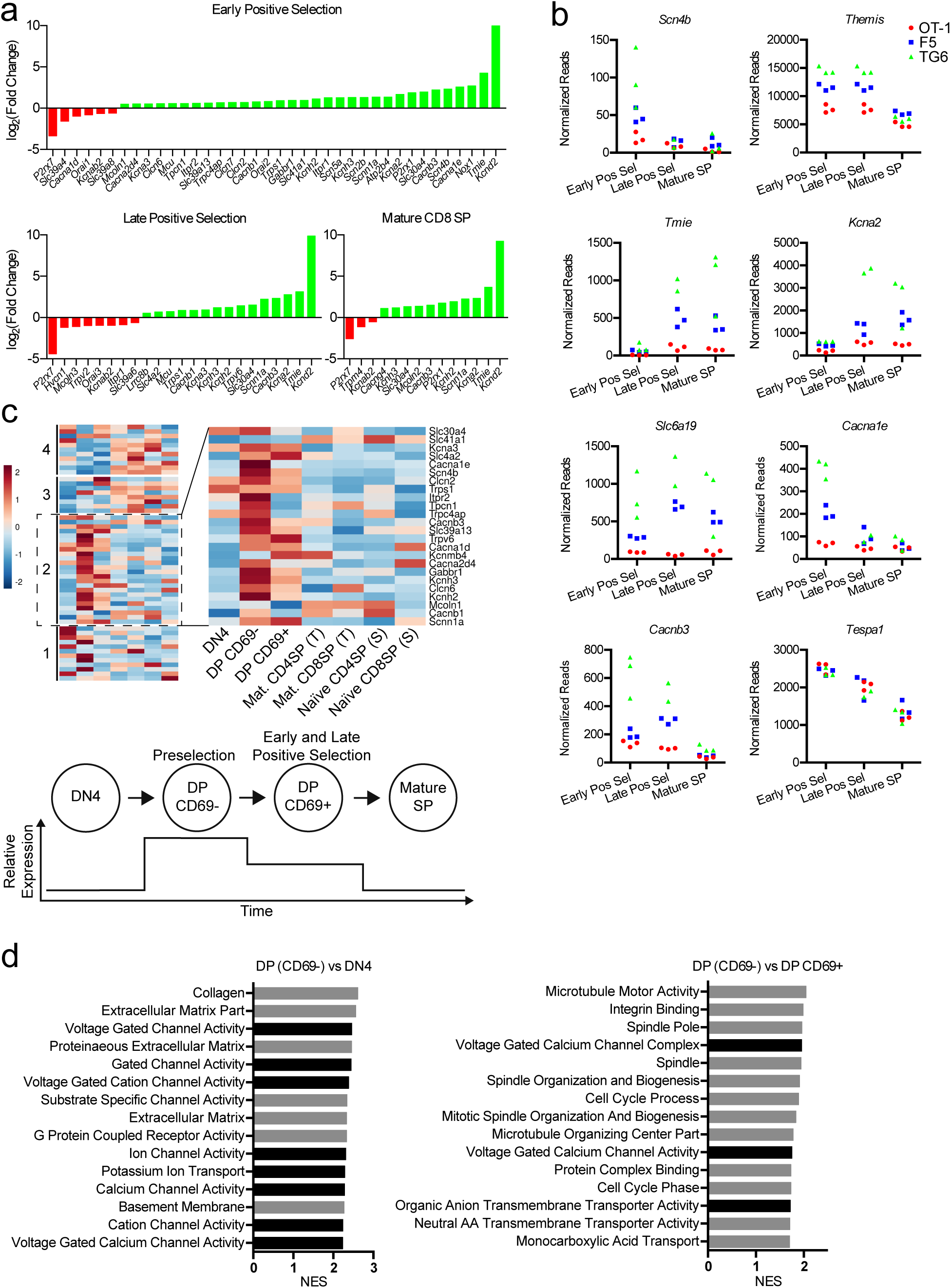
Thymocytes with low self-reactivity retain a preselection gene expression program marked by elevated expression of ion channel genes. (a) Ion channel genes from the manually curated list in Figure 7b that exhibit significantly different (padj<0.05, fold change>0.5) expression between OT1 versus TG6 thymocytes at the indicated stages of development. Positive values (green) indicate genes enriched in TG6; negative values (red) indicate genes enriched in OT-1. (b) Normalized read counts (see methods) for the indicated genes at each stage of development. Red circles: OT-1; Blue squares: F5; Green triangles: TG6. (c) Left: Heatmap of normalized expression of the ion transport genes (Group 2 genes from Figure 7d) in wild thymocytes at the indicated stages of development (scaled per row). Data are from the ImmGen microarray database. Bottom: Graphic representation of the average gene expression pattern. (d) Normalized enrichment score (NES) for the top 15 gene sets enriched in DP CD69-cells compared to DN4 and DP CD69+ thymocytes, using gene set enrichment analysis (GSEA) and the Gene Ontology database. Black bars indicate gene sets relating to ion channels.

## Supplemental Tables

**Supplemental Table 1:** TG6 thymocytes spend less time signaling compared to OT-1 cells. Percent time signaling was calculated by dividing the number of signaling timepoints by total timepoints.

|  | **TG6**<br>**3h** | **OT-1**<br>**3h** | **TG6**<br>**6h** | **OT-1**<br>**6h** |
| --- | --- | --- | --- | --- |
| **Total Signaling Timepoints** | 172 | 409 | 177 | 163 |
| **Total Timepoints** | 16928 | 21973 | 32367 | 14553 |
| **Percent Time Signaling** | 1.02 | 1.86 | 0.55 | 1.12 |

**Supplemental Table 2:** TG6 thymocytes experience less frequent TCR signaling than OT-1 thymocytes. The frequency of signaling events was calculated by dividing the total number of signaling events by the cumulative track imaging time (the sum of all of the track durations for all runs) for each condition.

|  | **TG6 3h** | **OT-1 3h** | **TG6 6h** | **OT-1 6h** |
| --- | --- | --- | --- | --- |
| **Total Signaling Events** | 42 | 61 | 44 | 32 |
| **Cumulative Track Imaging Time (h)** | 47.022 | 61.036 | 89.908 | 40.425 |
| **Signaling Events/Hour** | 0.89 | 1.00 | 0.49 | 0.79 |

**Supplemental Table 3:**
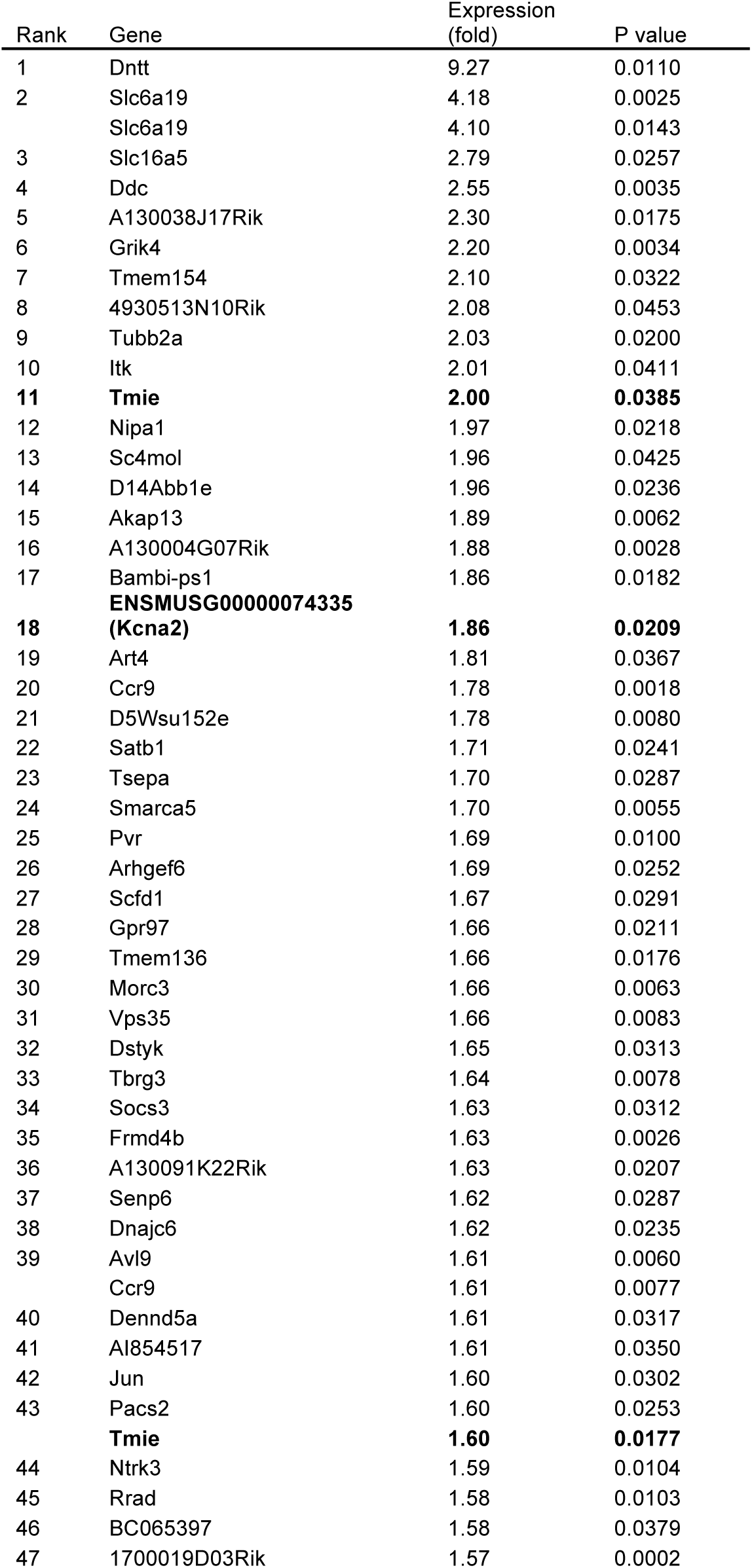
Genes upregulated in CD5lo compared to CD5hi naïve polyclonal CD8SP T cells by microarray analysis. The top 47 significantly different (P<0.05, >1.45 fold difference) genes upregulated in sorted CD5lo naive polyclonal CD8SP T cells, by microarray analysis, and ranked by their expression (fold) difference. Gene that appear multiple times represent multiple probe sets for the same gene, and a number in the left-most column is included only for the first listing. Genes of interest, Kcna2 and Tmie, are in bold. Data was analyzed by Fulton et al (GSE62142).

**Supplemental Table 4:** Genes upregulated in CD5lo compared to CD5hi naïve polyclonal CD8SP T cells by RNA-seq analysis. The top 51 significantly different (padj<0.05) upregulated genes in sorted CD5lo naive polyclonal CD8+ T cells and ranked by their expression (fold) difference, from the Matson, et al. RNA-seq dataset (GSE151395). Genes of interest, *Kcna2* and *Tmie*, are in bold. Data was analyzed by DEseq2.

| Rank | Gene | Expression (fold) | padj |
| --- | --- | --- | --- |
| 1 | Col2a1 | 5.95 | 7.25E-05 |
| 2 | Irs1 | 5.91 | 6.68E-04 |
| 3 | Meg3 | 5.87 | 2.58E-02 |
| 4 | Col9a2 | 4.81 | 1.13E-02 |
| 5 | Igf2 | 4.28 | 1.17E-03 |
| 6 | Lect1 | 4.25 | 1.06E-02 |
| 7 | P3h3 | 4.00 | 3.99E-02 |
| 8 | Mir6934 | 3.62 | 2.65E-02 |
| 9 | Thbs1 | 3.61 | 3.01E-02 |
| 10 | Pcsk1 | 3.52 | 2.74E-07 |
| 11 | 5930430L01Rik | 3.35 | 4.95E-05 |
| 12 | Slc6a19 | 3.20 | 3.39E-14 |
| 13 | Col12a1 | 3.12 | 1.29E-02 |
| 14 | Grm6 | 3.00 | 6.58E-10 |
| 15 | Dntt | 2.94 | 3.83E-21 |
| 16 | Lrrc3b | 2.93 | 1.08E-05 |
| 17 | Adamts14 | 2.67 | 7.45E-21 |
| 18 | Tdrd5 | 2.55 | 5.10E-06 |
| 19 | Zfp36l3 | 2.52 | 4.89E-02 |
| 20 | Nrg2 | 2.30 | 2.22E-03 |
| 21 | Fbxl21 | 2.22 | 2.17E-04 |
| 22 | Kcnj16 | 2.20 | 4.85E-06 |
| 23 | Btc | 2.17 | 1.60E-02 |
| 24 | Islr | 2.14 | 5.34E-12 |
| 25 | Slc6a19os | 2.09 | 7.51E-07 |
| 26 | P4ha2 | 2.01 | 5.01E-09 |
| 27 | Nphs2 | 1.98 | 7.03E-04 |
| 28 | A130077B15Rik | 1.96 | 4.63E-06 |
| 29 | 4930447K03Rik | 1.94 | 2.69E-02 |
| 30 | Fbln2 | 1.92 | 4.58E-05 |
| 31 | Arpp21 | 1.91 | 6.49E-07 |
| 32 | Osgin1 | 1.89 | 1.61E-09 |
| 33 | Gpr34 | 1.89 | 1.50E-05 |
| 34 | 4930512J16Rik | 1.82 | 4.43E-02 |
| 35 | Pomgnt2 | 1.81 | 2.95E-03 |
| 36 | Slc16a5 | 1.72 | 5.13E-29 |
| 37 | Cyp2s1 | 1.70 | 1.19E-13 |
| 38 | Dmbt1 | 1.65 | 1.76E-18 |
| 39 | Slc6a9 | 1.61 | 1.41E-02 |
| 40 | Gm10516 | 1.61 | 1.84E-02 |
| 41 | Prdm1 | 1.53 | 9.84E-08 |
| 42 | Prss50 | 1.52 | 3.91E-03 |
| 43 | Ntn5 | 1.51 | 4.42E-02 |
| 44 | Rmrp | 1.48 | 2.13E-03 |
| **45** | **Tmie** | **1.47** | **4.08E-04** |
| 46 | Igsf23 | 1.47 | 4.61E-15 |
| 47 | Smyd1 | 1.42 | 1.03E-03 |
| 48 | Zp3r | 1.41 | 1.81E-02 |
| 49 | Angptl2 | 1.41 | 3.47E-06 |
| 50 | Bmf | 1.38 | 3.00E-07 |
| **51** | **Kcna2** | **1.35** | **1.92E-13** |

## Supplemental Movies

(please see Supporting Files)

<u>Movie S1: Representative TG6 thymocyte signaling event.</u> Preselection TG6 thymocytes were loaded with the ratiometric calcium indicator dye Indo1LR and allowed to migrate into selecting thymic slices for 6hrs, then imaged using two-photon-microscopy. Video is a z-projection of the ratio of fluorescent signal in the calcium-bound channel over calcium-unbound channel for Indo1LR displayed as a heatmap (red=calcium high, purple=calcium low). Arrow indicates timepoints in which the cell displays elevated calcium (130 seconds). Frames were collected every 10s for 10.3min. Dimensions: 45.7μm (width) by 75.0μm (height) by 24.9μm (depth). Scale bar is 10μm.

<u>Movie S2: Representative OT-1 thymocyte signaling event.</u> Preselection OT-1 thymocytes were loaded with the ratiometric calcium indicator dye Indo1LR and allowed to migrate into selecting thymic slices for 6hrs, then imaged using two-photon-microscopy. Video is a z-projection of the ratio of fluorescent signal in the calcium-bound channel over calcium-unbound channel for Indo1LR displayed as a heatmap (red=calcium high, purple=calcium low). Arrow indicates timepoints in which the cell displays elevated calcium (230 seconds). Frames were collected every 10s for 14.3min. Dimensions: 68.9μm (width) by 49.8μm (height) by 18μm (depth). Scale bar is 10μm.

